# Transcription initiation mapping in 31 bovine tissues reveals complex promoter activity, pervasive transcription, and tissue-specific promoter usage

**DOI:** 10.1101/2020.09.05.284547

**Authors:** D.E. Goszczynski, M.M. Halstead, A.D. Islas-Trejo, H. Zhou, P.J. Ross

**Affiliations:** Department of Animal Science, University of California, Davis, CA, USA 95616

**Author notes:** Corresponding author: Ross PJ. Both authors contributed equally to this work (D.E. Goszczynski, M.M. Halstead).

**Keywords:** transcription start sites, RAMPAGE, cattle, promoter, atlas

## Abstract

Characterizing transcription start sites is essential for understanding the regulatory mechanisms that control gene expression. Recently, a new bovine genome assembly (ARS-UCD1.2) with high continuity, accuracy, and completeness was released; however, the functional annotation of the bovine genome lacks precise transcription start sites and contains a low number of transcripts in comparison to human and mouse. Using the RAMPAGE approach, this study identified transcription start sites at high resolution in a large collection of bovine tissues. We found several known and novel transcription start sites attributed to promoters of protein coding and lncRNA genes that were validated through experimental and *in silico* evidence. With these findings, the annotation of transcription start sites in cattle reached a level comparable to the mouse and human genome annotations. In addition, we identified and characterized transcription start sites for antisense transcripts derived from bidirectional promoters, potential lncRNAs, mRNAs, and pre-miRNAs. We also analyzed the quantitative aspects of RAMPAGE to produce a promoter activity atlas, reaching highly reproducible results comparable to traditional RNA-seq. Co-expression networks revealed considerable use of tissue specific promoters, especially between brain and testicle, which expressed several genes in common from alternate loci. Furthermore, regions surrounding co-expressed modules were enriched in binding factor motifs representative of each tissue. The comprehensive annotation of promoters in such a large collection of tissues will substantially contribute to our understanding of gene expression in cattle and other mammalian species, shortening the gap between genotypes and phenotypes.

## INTRODUCTION

With the advent of new sequencing technologies, genomics has emerged as one of the most promising fields in biology. Since its implementation in breeding schemes, livestock industries have seen enormous improvements in productivity (Van Eenennaam et al. 2014), and genomics are expected to play a key role in improving animal welfare, protecting the environment, and securing high quality food for a growing human population (Britt et al. 2018). For genomic approaches to succeed, high quality reference genomes are essential. The first bovine genome was published in 2009 (Bovine Genome Sequencing and Analysis Consortium 2009). Its publication and the improvements that followed have allowed the application of genomic technologies aimed at genetic improvement (Null et al. 2019), the study of molecular and physiological mechanisms of animal health and disease (Ju et al. 2020), and the development of genome engineering approaches (Bevacqua et al. 2016). Recently, a new version of the bovine genome, ARS-UCD1.2, was assembled using single molecule sequencing, increasing the continuity, accuracy, and completeness compared to its predecessor (Rosen et al. 2020). With a more reliable reference genome available, the focus now transitions from genome sequence to genome features. The annotation of functional regulatory elements in cattle, as well as other important livestock species, is one of the main objectives of the international Functional Annotation of Animal Genomes (FAANG) Consortium (www.faang.org) (Andersson et al. 2015).

A key step to study transcription regulation is the elucidation of transcription start sites (TSSs), as these loci harbor transcription factor binding sites that drive gene expression and integrate the inputs from other cis-regulatory elements (e.g., enhancers). Generally, TSSs are amidst an epigenetic landscape that involves chromatin accessibility (Meers et al. 2018), H3K4me3 and H3K27ac histone marks (Ernst and Kellis 2017). These epigenetic characteristics can be used to differentiate promoters from other gene regulatory regions, such as enhancers, which are typically enriched in additional histone marks such as H3K4me1 (Ernst and Kellis 2017). Methods to identify 5’ ends of transcripts have been based on the capture of 5’ caps or the preferential ligation of adapters to them (Batut et al. 2013; Kodzius et al. 2006; Salimullah et al. 2011; Yamashita et al. 2011), as opposed to RNA-seq methods, where 5’ ends are frequently underrepresented due to premature termination of reverse transcription (RT), especially when oligo(dT) primers are used. The RAMPAGE technique (RNA Annotation and Mapping of Promoters for the Analysis of Gene Expression) is an accurate 5’-complete cDNA sequencing approach that allows for *ab initio* identification of TSSs at single base pair resolution and quantification of their expression (Batut et al. 2013). Briefly, RAMPAGE evaluates statistical enrichment (peaks) of 5’ signal across the genome and merges peaks in close proximity to each other into transcription start site clusters (TSCs). It is worth mentioning that the TSC term was used in the original RAMPAGE publication to refer to the pipeline output, but TSCs are conceptually equivalent to TSSs; in this manuscript, we will maintain the use of the original nomenclature. In comparison to other protocols, RAMPAGE is simple, scalable, requires low inputs, allows for discrimination of nonspecific signal caused by incomplete cDNAs and of PCR duplicates (Batut et al. 2013).

Considering the start coordinate of an annotated transcript as the transcription start site (TSS), the ARS-UCD1.2 bovine genome annotation from the Ensembl database (release 95) includes 39,705 TSSs for protein coding or lncRNA transcripts: a low number in comparison to the 139,158 TSSs annotated in human (GRCh38) and the 81,573 TSSs annotated in mouse (GRCm38). Even though small systematic differences are expected in the number of promoters between species, experimental evidence indicates that several bovine transcripts remain to be characterized and annotated (Tong et al. 2017; Weikard et al. 2013). Moreover, studies in mouse and human have demonstrated that a significant proportion of TSSs are enriched in evolutionarily conserved regions (Abugessaisa et al. 2019; FANTOM Consortium and the RIKEN PMI and CLST (DGT) 2014), suggesting the number of TSSs in the bovine genome should be higher.

As part of the FAANG Consortium, our main objective was to contribute to the bovine transcriptome annotation by identifying TSCs in a large collection of tissues using the RAMPAGE approach. Identified TSCs were validated through several strategies that included experimental evidence from chromatin accessibility assays (ATAC-seq) and histone modification profiling (ChIP-seq), which included H3K4me1, H3K4me3, H3K27ac, and H3K27me3. In addition, we evaluated sequence enrichment for typical promoter elements such as TATA, GC, and CCAAT boxes in regions surrounding TSCs, and known tissue-specific transcription factor binding motifs. The RAMPAGE signal was implemented into a promoter activity atlas that contemplates transcription initiation in each tissue. To evaluate the quantitative aspect of the technique, we compared it to conventional RNA-seq, which revealed highly consistent and reproducible expression quantification. Unassigned RAMPAGE peaks with insufficient evidence for gene assignment were screened through a set of bioinformatic tools to evaluate their potential as unannotated mRNAs, pre-miRNAs, and ncRNAs. Some of these peaks were particularly interesting due to their antisense activity. Lastly, expression values derived from the RAMPAGE signal were used to generate co-expression networks and to characterize the use of alternative TSCs across tissues.

## RESULTS

### Dataset complexity

A total of 111 biological samples corresponding to 31 different tissues from two male (M) and two female (F) individuals were sequenced in nine batches, resulting in 2.72 billion fragments. Most fragments from each library mapped uniquely to the reference genome, yielding a total of 2.39 billion (88%) uniquely mapped fragments (Supplemental Table S1). After duplicate removal, the number of fragments decreased to 673 million (29% of uniquely mapped), with an average of 6.1 million (SD 3.8 million) fragments per sample (Supplemental Fig S1A). A preliminary evaluation by hierarchical clustering based on gene expression (independent of peak calling) allowed us to identify seven outliers: abomasum-M1, bladder-F2, colon-F1, esophagus-M1, esophagus-M2, lung-M1, and trachea-M1 (Supplemental Fig S1B). All outliers were discarded from further analyses. Noise level, measured as the proportion of read 1s outside RAMPAGE peaks, was below 20%.

**Supplemental Table S1.** RAMPAGE sequencing summary. The first sheet contains the number of sequenced fragments and alignment rates for each sample in each sequencing run. The second sheet contains the number of uniquely mapped reads depleted of duplicates, which were used for peak calling after combining sequences from each sample.

**Supplemental Fig S1.**
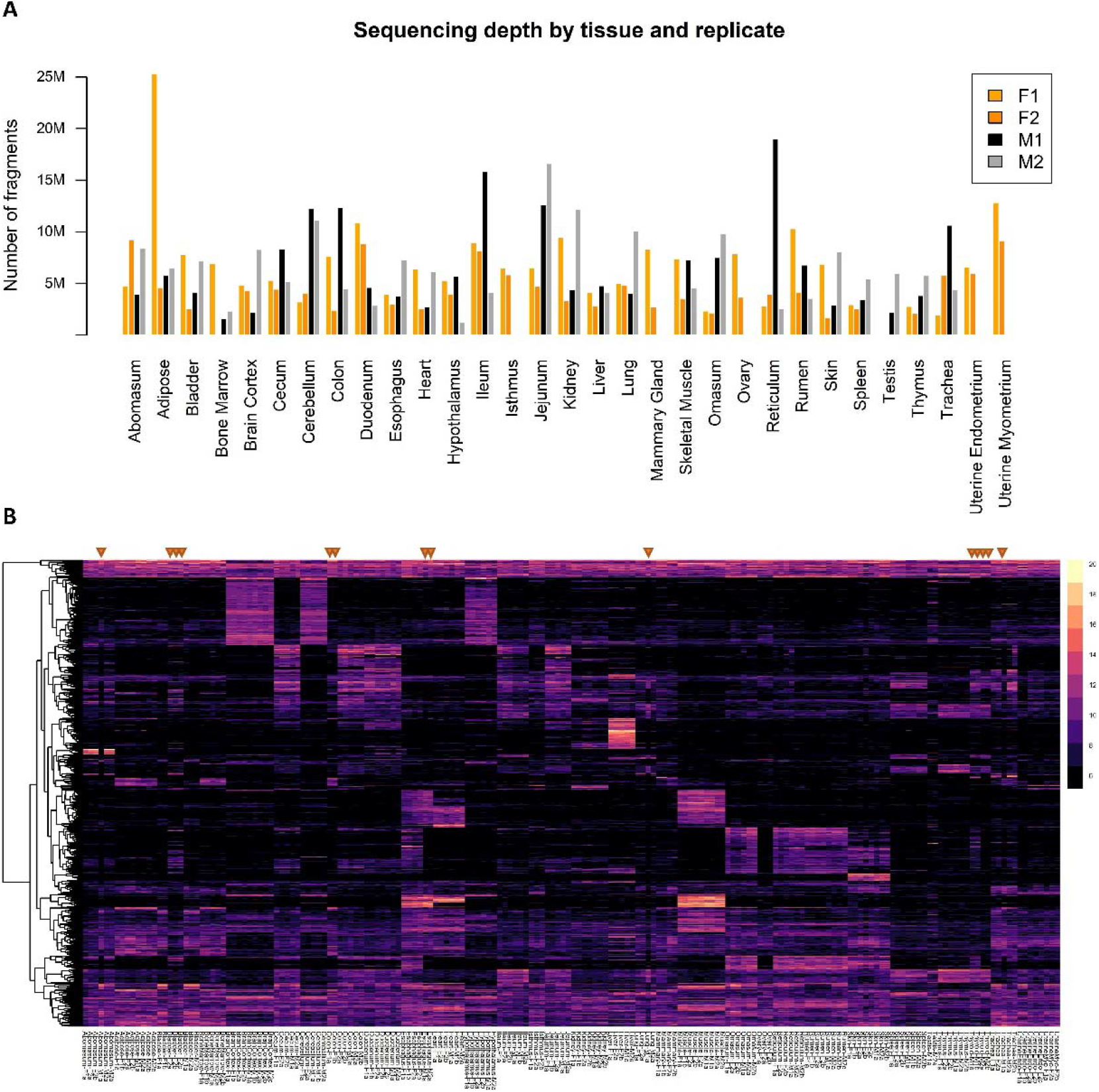
Dataset composition. (A) Sequencing depth by tissue and replicate. The y axis indicates the number of uniquely mapped fragments after duplicate removal. (B) Profiling by gene expression (variance-stabilized counts) showing the unclear identity of the abomasum-M1, bladder-F2, colon-F1, esophagus-M1, esophagus-M2, lung-M1, and trachea-M1 samples (indicated with orange triangles), which were excluded from further analyses.

### Identification of TSCs by RAMPAGE

The analysis of 104 samples from 31 tissues from four individuals (two males, two females) identified 93,767 TSCs between the combined dataset (87,553) and single tissue datasets (6,214). TSC size varied from a single base to intervals as broad as 1,089 bp, with most elements falling in the range of 5 to 25 bp (Figure 1A). TSCs identified in the combined dataset were supported by a minimum of 22 independent RAMPAGE tags (Figure 1B), as opposed to tissue specific TSCs, which were supported by a minimum of three RAMPAGE tags but were conserved between biological replicates.

**Figure 1.**
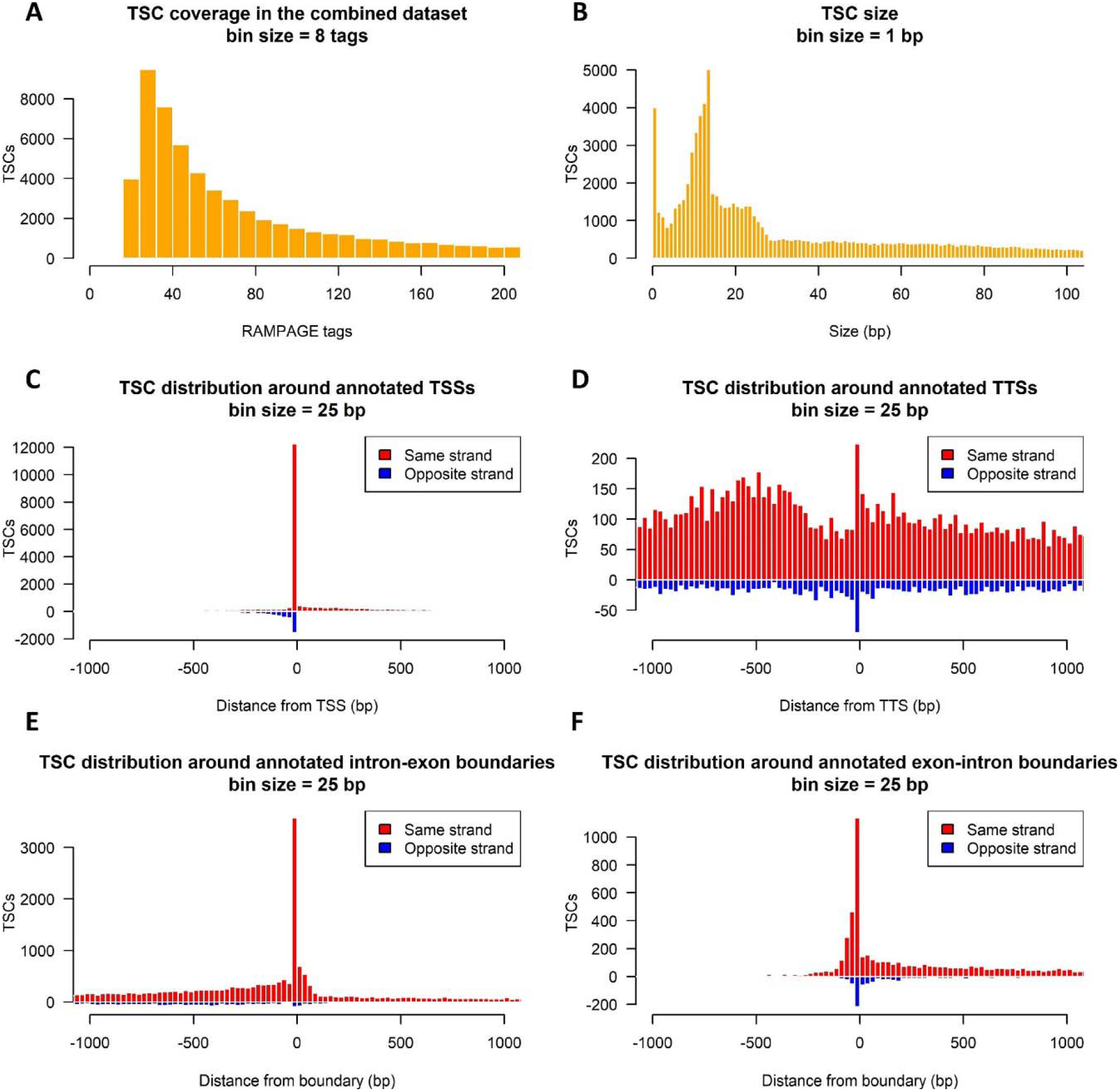
Identification of promoters by RAMPAGE. (A) Tag coverage in TSCs identified in the combined dataset. (B) TSC size distribution for the whole dataset. (C) Histogram of the distance between TSCs and their nearest annotated TSS. (D) Histogram of the distance between TSC and the nearest annotated TTS. (E) Histogram of the distance between TSCs and intron-exon boundaries. Only intron-exon boundaries not overlapping with any annotated TSSs were considered. (F) Histogram of the distance between TSCs and exon-intron boundaries. For visualization, exon-intron boundaries corresponding to the end of the first exon were ignored, as TSCs tended to group immediately upstream of these boundaries and interfere with the visualization.

The bovine genome annotation (Ensembl release 95) considered for this study comprised 33,019 and 31,052 different TSSs and transcription termination sites (TTSs), respectively, associated with 23,328 protein coding or long noncoding (lncRNA) genes. To account for nonspecific transcription initiation, TSSs were defined as annotated start coordinate ± 50 bp. The annotation contained 99 TSSs overlapping with TTSs, 1,676 TSSs overlapping with intron-exon boundaries, and 850 TSSs overlapping with exon-intron boundaries. It is worth mentioning that 662 of the latter overlapped with boundaries at the end of the first exon due to short exon length.

Most of the TSCs identified in this study were absent from the Ensembl annotation. In fact, only 12,072 (13%) of the TSCs overlapped with previously annotated TSSs (Figure 1C). In order to determine whether the remaining 19,781 annotated TSSs were undetected or shifted with respect to our findings, we first evaluated their gene expression and, when their expression level was higher than 3 counts per million (CPM), we mapped their location relative to RAMPAGE TSCs. No expression was detected for 3,620 (16%) of the annotated genes, which corresponded to 3,718 annotated TSSs. The location of the remaining 16,063 nonoverlapped TSSs around TSCs suggested that 44% of these elements were shifted by a margin of 200 bp (Supplemental Fig S2). No TSC enrichment was observed in the vicinity of TTSs (Figure 1D). On the other hand, several TSCs were found to overlap with internal annotated intron-exon (2,148) (Figure 1E) and exon-intron boundaries (299) by at least one base (Figure 1F).

To evaluate the possibility that boundary associated TSCs surged as a consequence of transcript fragmentation or capture of partial splicing products, we investigated chromatin accessibility and histone mark profiles in tissues where TSCs were active. This analysis revealed the presence of transcriptional activation marks (H3K4me3, H3K27ac) and accessible chromatin (ATAC-seq) at the TSCs, while control intron-exon and exon-intron boundaries showed no enrichment for these marks (Supplemental Fig S3).

**Supplemental Fig S2.**
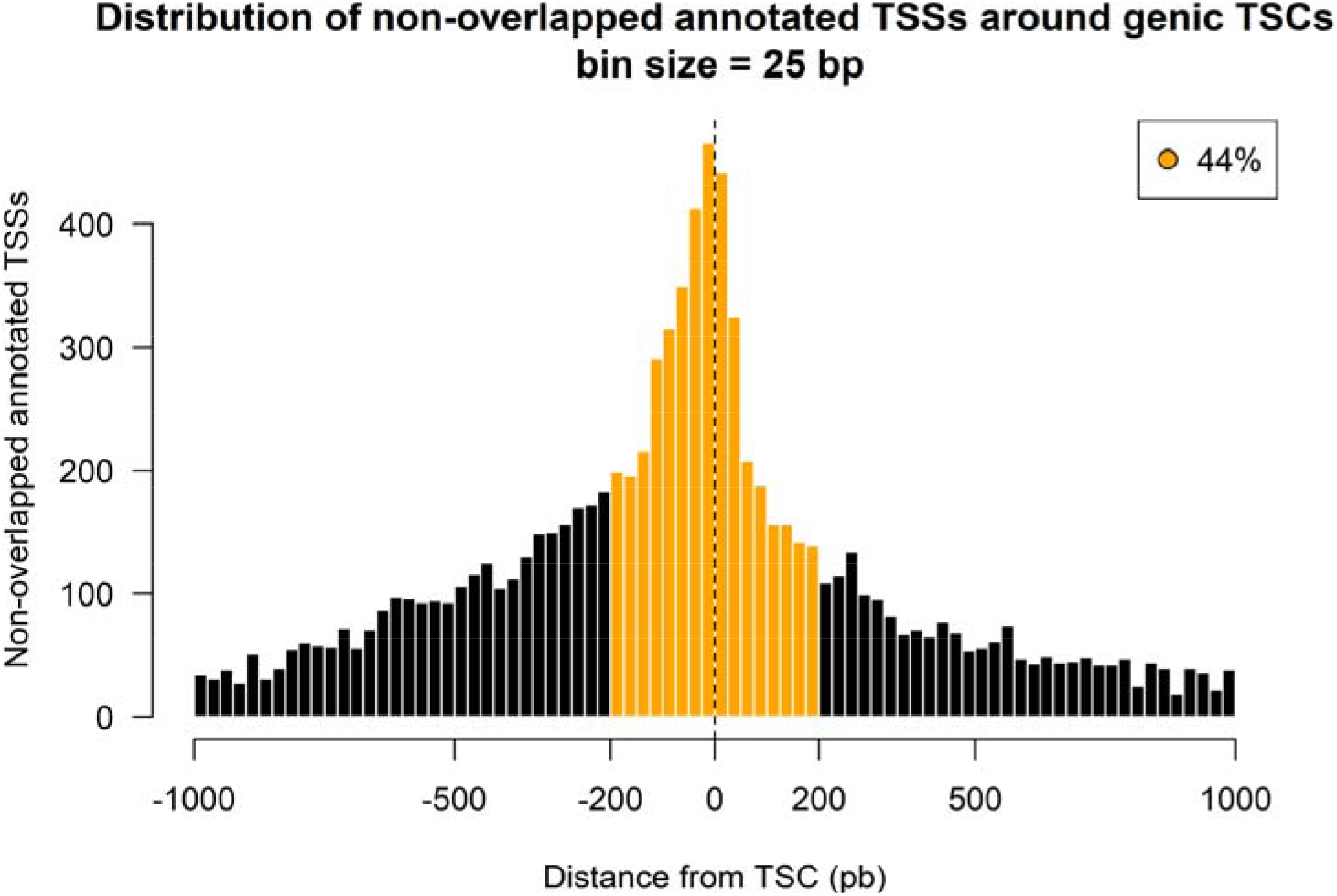
Distribution of nonoverlapped annotated TSSs relative to genic TSCs identified by RAMPAGE. Many of the annotated TSSs were shifted with respect to TSCs by a margin of 200 bp (orange), while the rest likely belonged to transcripts variants undetected in our dataset (black). Only annotated genes with high expression levels (CPM > 3) were considered for this figure.

**Supplemental Fig S3.**
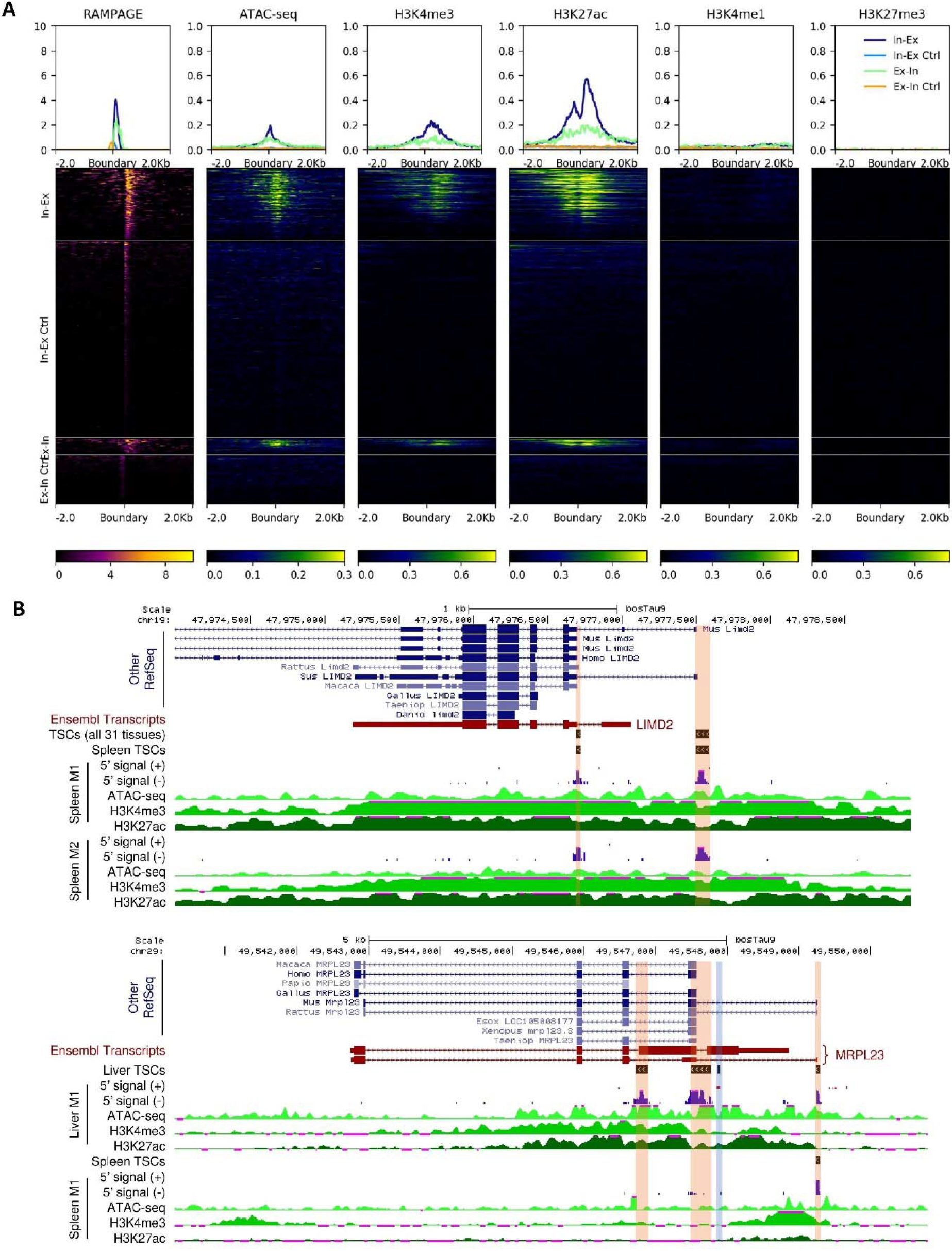
Chromatin accessibility and histone marks at intron-exon and exon-intron boundaries associated with TSCs (Lung-M2). (A) Distribution of reads from different assays over a window of 2 kb to each side of the boundaries. The RAMPAGE signal contemplates only the first read of the pair (R1). Panels ‘In-Ex’ and ‘Ex-In’ represent intron-exon and exon-intron boundaries overlapped by TSCs and expressed at a minimum level of 3 CPM with respect to all RAMPAGE tags. Control regions (Ctrl) comprised boundaries from the same transcripts, covered by at least 3 fragments, and located at least 2 kb away from the nearest TSC to avoid signals from near TSCs. Heatmaps are colored according to CPM values. Profiles above heatmaps represent the median signal at each coordinate of the 4kb window. The co-occurrence of chromatin accessibility (ATAC-seq) and transcriptional activation marks (H3K4me3, H3K27ac), and the absence of enhancer (H3K4me1) and repressive (H3K27me3) marks at the TSC sites implicated these boundaries as putative promoters. (B) 5’ tags and epigenetic signals at boundary-associated TSCs. We found no evidence supporting the annotated TSS for the bovine *LIMD2* gene. In addition, one of the TSCs overlapped an annotated intron-exon boundary. In case of the *MRPL23* gene, we found a highly active TSC at an exon-intron boundary, a second TSC between an exon-intron and an intron-exon boundary, and a third TSC supporting one of the annotated TSSs.

### Assignment of TSCs to genes by sequence evidence

Identified TSCs were assigned to genes based on the alignments of their fragments. We associated 46,708 ‘genic’ TSCs with 18,312 protein coding and lncRNA genes, accounting for 78% of the 23,328 annotated genes. In addition to these highly associated TSCs, we identified 3,789 elements weakly associated with 3,176 genes (< 3 reads), 308 of which were absent from the previous list. Undetected genes were enriched in GO terms associated with olfaction (Padj = 2.6E-165), taste (Padj = 1.3E-7), phototransduction (Padj = 9.5E-2), and defensins (Padj = 1.0E-4). In addition, undetected genes were strongly enriched in homeobox terms (Padj = 3.9E-18), which evidenced the underrepresentation of embryonic and fetal genes in the dataset.

The number of genes detected in each biological sample (CPM of genic TSC > 3) varied from 7,841 (muscle-M1) to 11,281 (trachea–M2), with an average of 10,009 genes per sample. Based on the Ensembl annotation, 36% of the genic TSCs were located <2.5 kb upstream of annotated TSSs, 31% were in exons, 23% were in introns, and 10% were in intergenic loci (>2.5 kb from annotated TSS) (Figure 2A). Similar to the global set of TSCs, the size distribution of genic TSCs showed a narrow peak at 1 bp and a broad peak between 5 and 25 bp (Figure 2B).

**Figure 2.**
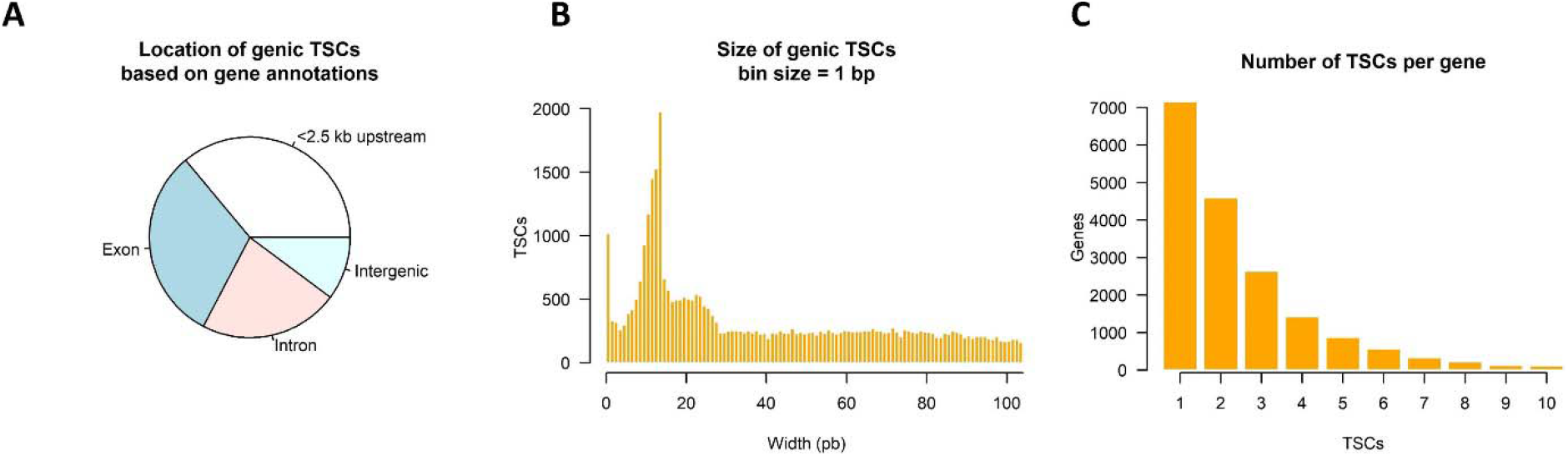
Genic TSCs identified by RAMPAGE sequencing. (A) Location of genic TSCs according to gene annotations. (B) Histogram of TSC size. (C) Use of multiple TSCs for the same gene. Genes with more than ten TSCs were excluded from the plot as they were likely affected by technical artifacts.

The assignment of TSCs to genes revealed that 61% of the detected genes were transcribed from more than one site (Figure 2C). We found an average of 2.6 TSC/gene, which was twice the value calculated from the Ensembl annotation (1.3 TSS/gene). Comparatively, our TSC/gene value was between the GRCm38 mouse annotation (2.3 TSS/gene) and the GRCh38 human annotation (2.9 TSS/gene). It is worth mentioning that since RAMPAGE TSCs were identified by merging peaks within windows of 150 bp, we merged annotated TSSs within 150 bp to make comparisons fair.

### Identification of novel promoters

As mentioned previously, most of the genic transcripts in our dataset (34,649) originated from TSCs that were hundreds or thousands of base pairs away from the nearest annotated TSS. To characterize and validate these ‘novel’ genic TSCs, we studied their location based on gene annotations and their distance to genic TSCs also found in the annotation. Then, we evaluated the presence of typical promoter binding motifs and epigenetic marks for transcriptional activation in regions surrounding TSCs.

According to gene annotations, most of the novel genic TSCs were in exons (38%), followed by introns (30%), regions less than 2.5 kb upstream of annotated TSSs (18%), and intergenic regions (14%) (Figure 3A). Novel genic TSCs were abundant within 1 kb upstream and 2 kb downstream of annotated TSSs (Figure 3B), constituting new alternative sites for transcription initiation in such genes. In terms of expression, novel TSCs contributed substantially to the transcriptome, with half of them driving more than 20% of the total gene expression (Figure 3C).

**Figure 3.**
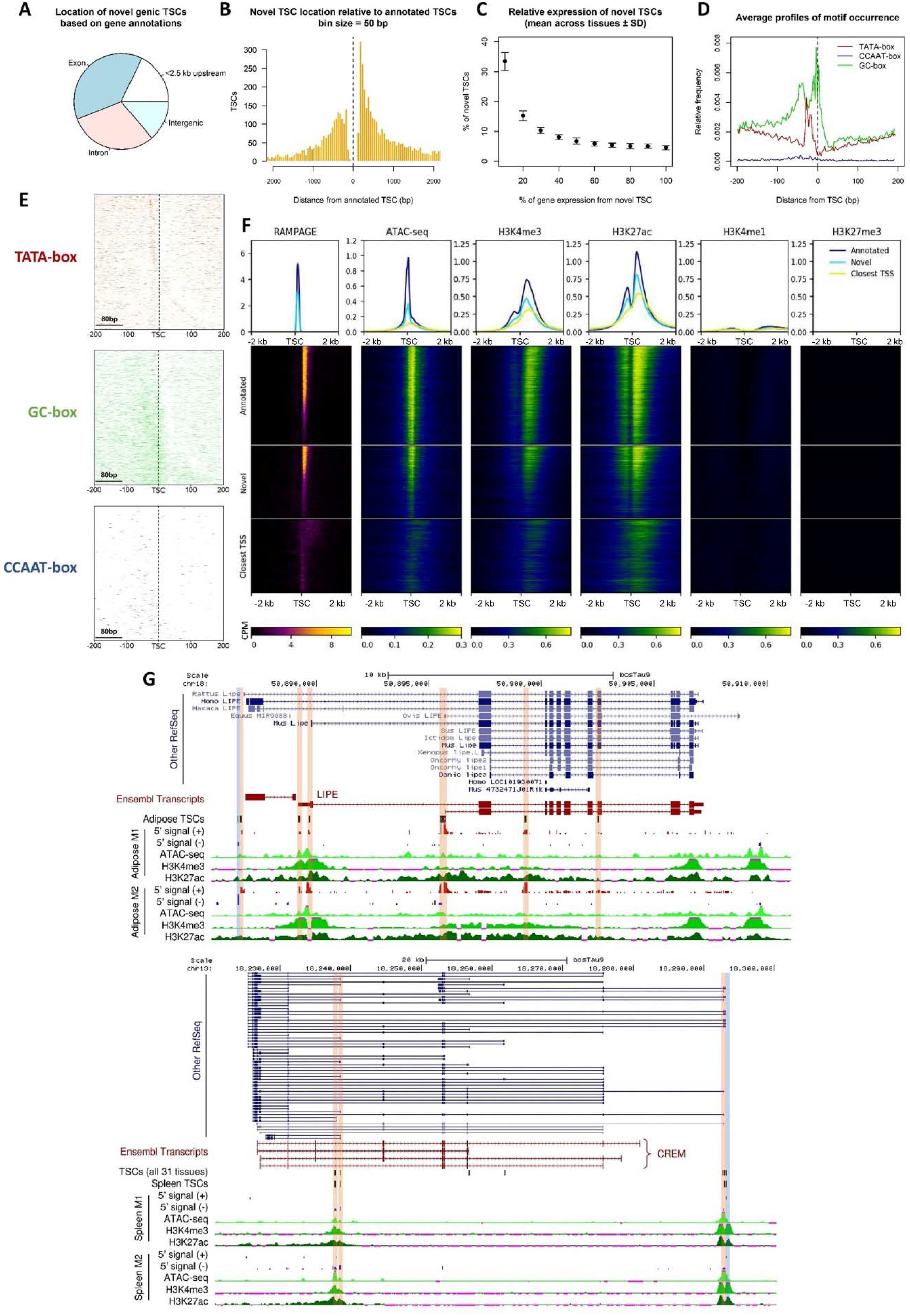
Novel promoters identified by RAMPAGE. (A) Location of novel TSCs according to gene annotations. (B) Histogram of the distance between novel TSCs and annotated TSCs. (C) Relative expression of novel TSCs. (D) Profiles of motif occurrence for TATA, CCAAT and GC boxes around novel TSCs (± 200 bp). (E) Motif density maps for TATA, CCAAT and GC-boxes around novel TSCs (± 200 bp). TSCs are ordered ascendingly by size, i.e., upper rows represent narrow TSCs. The zero coordinate represents the 5’ end of the TSC. Narrow TSCs were particularly enriched with TATA-box motifs and broad TSCs were enriched with GC-box motifs. (F) Epigenetic marks at reported and novel TSCs. The co-occurrence of chromatin accessibility and transcriptional activation marks (H3K4me3, H3K27ac) at novel TSCs, as well as the absence of poised enhancer and repressive marks (H3K4me1, H3K27me3), suggested these TSCs constituted promoter regions. The fuzzy signal observed around the closest annotated TSSs evidenced the absence of annotation for these novel elements. The data shown in the figure corresponds to the lung-M2 sample (>3 CPM). Heatmaps are colored according to CPM values. (G) Novel TSCs for the *LIPE* and *CREM* genes. Most of the new variants are supported by annotations from other species. Antisense TSCs were marked with blue to distinguish them from sense TSCs (orange).

We then evaluated the characteristics of sequences surrounding novel TSCs (Figure 3D). We found high abundance of TATA-boxes 25 bp upstream of novel TSCs, especially for narrow TSCs (Figure 3E). In general, broad TSCs included more than one TSS, explaining the presence of additional TATA-boxes downstream of the zero coordinate. GC-boxes were highly abundant in broad TSCs and localized mainly at the TSC site and 40 bp upstream. On the other hand, the abundance of CCAAT boxes was relatively low compared to the other marks, but when present, they localized 10-70 bp upstream of broad TSCs. Binding motif enrichment showed these three motifs were significantly enriched in regions 300 bp upstream to 100 bp downstream of novel TSCs compared to other sequences from the genome (p-value ^TATA^ = 1E-61, p-value ^CCAAT^ = 1E-178, p-value ^GC^ = 1E-183) as well as regions surrounding internal exons (300 bp upstream to 100 bp downstream; p-value ^TATA^ = 1E-14, p-value ^CCAAT^ = 1E-14, p-value GC = 1E-5). This data indicated that regions immediately upstream of novel genic TSCs have typical promoter characteristics, validating their identity as TSSs.

Evaluating the chromatin landscape surrounding TSCs, we observed a consistent co-occurrence of enriched chromatin accessibility, H3K4me3, and H3K27ac around novel genic TSCs, indicating these TSCs were associated with an epigenetic signature characteristic of transcriptional activation (Figure 3F). On the other hand, we detected low signal for H3K4me1 (enhancers) and no signal for H3K27me3 (silencing), which are typically found in enhancers and transcriptionally silent promoters, respectively. Additionally, we observed that the 5’ signal at the closest annotated TSS was generally lower and hundreds of base pairs away from the RAMPAGE TSCs (Figure 3F). Complementarily, we analyzed these epigenetic marks in subgroups of novel genic TSCs classified according to their genic location (exon, intron, intergenic, and promoters), which showed chromatin accessibility, H3K4me3, and H3K27ac enrichment in all subgroups (Supplemental Fig S4). Overall, the analyzed epigenetic information obtained from chromatin accessibility assays (ATAC-seq) and chromatin immunoprecipitation assays (ChIP-seq) from twelve of the analyzed biological samples, indicated that the novel TSCs associated with known genes represented functional TSSs (Figure 3G).

Some of the novel TSCs in exons demonstrated low epigenetic signal, especially for H3K4me3, suggesting they could represent artifacts produced by RNA degradation or incomplete cDNA synthesis. By generating histograms of the epigenetic signal at these TSCs, we noted a bimodal distribution that separated TSCs into groups with high and low epigenetic signal. By calculating the local minimum of signal, we estimated that approximately 30% of the novel TSCs in exons were in the latter group and could be potential artifacts (Supplemental Fig S5). These TSCs were labeled as such in Supplemental Table S2 (see below).

**Supplemental Fig S4.**
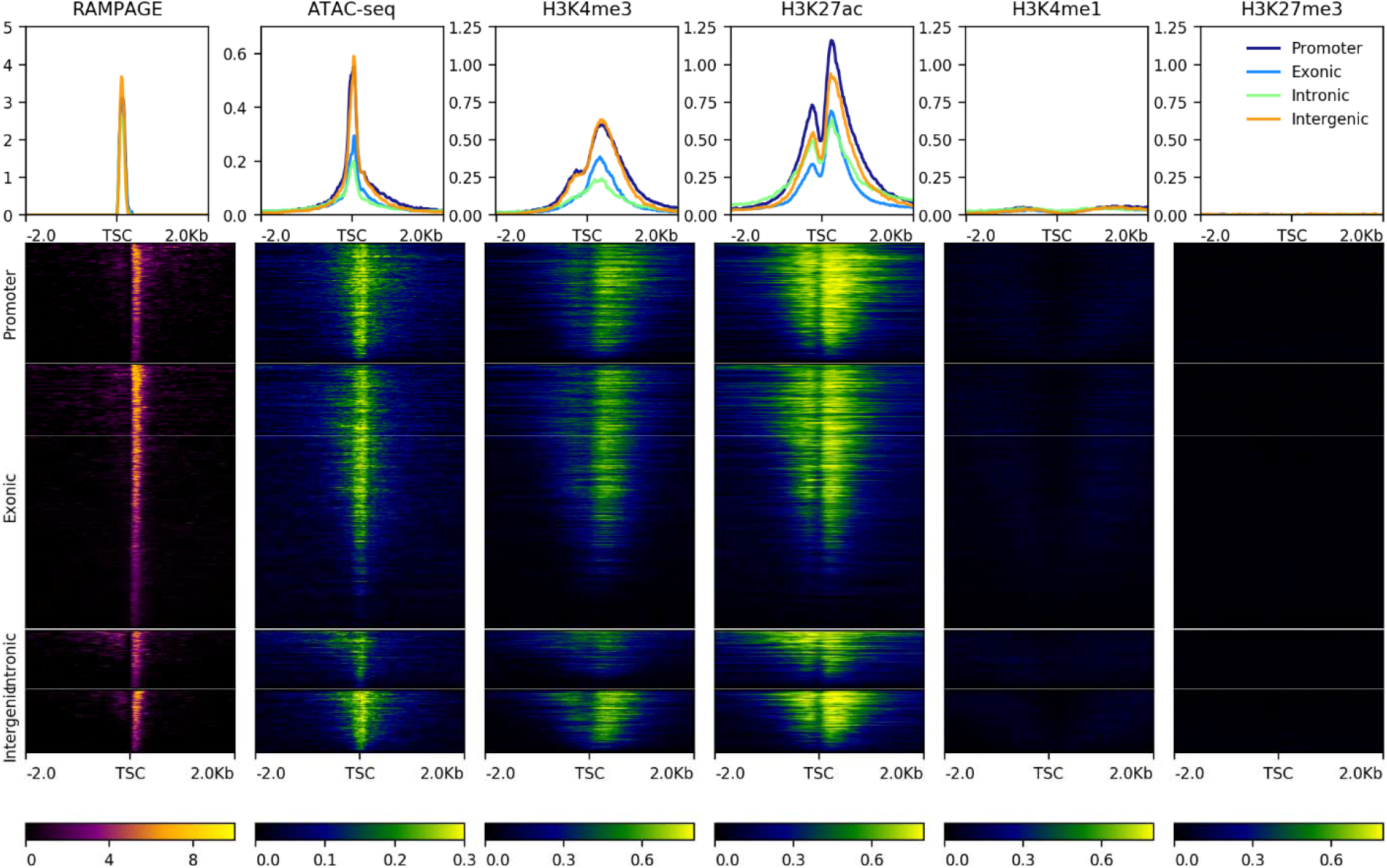
Epigenetic marks at subgroups of novel genic TSCs defined by their gene location (exon, intron, intergenic region, upstream of annotated TSS (promoter)) in the lung-M2 sample. A consistent co-occurrence of chromatin accessibility and transcriptional activation marks, as well as absence of poised enhancer and repressive marks, was observed in all subgroups. Heatmaps are colored according to CPM values.

**Supplemental Fig S5.**
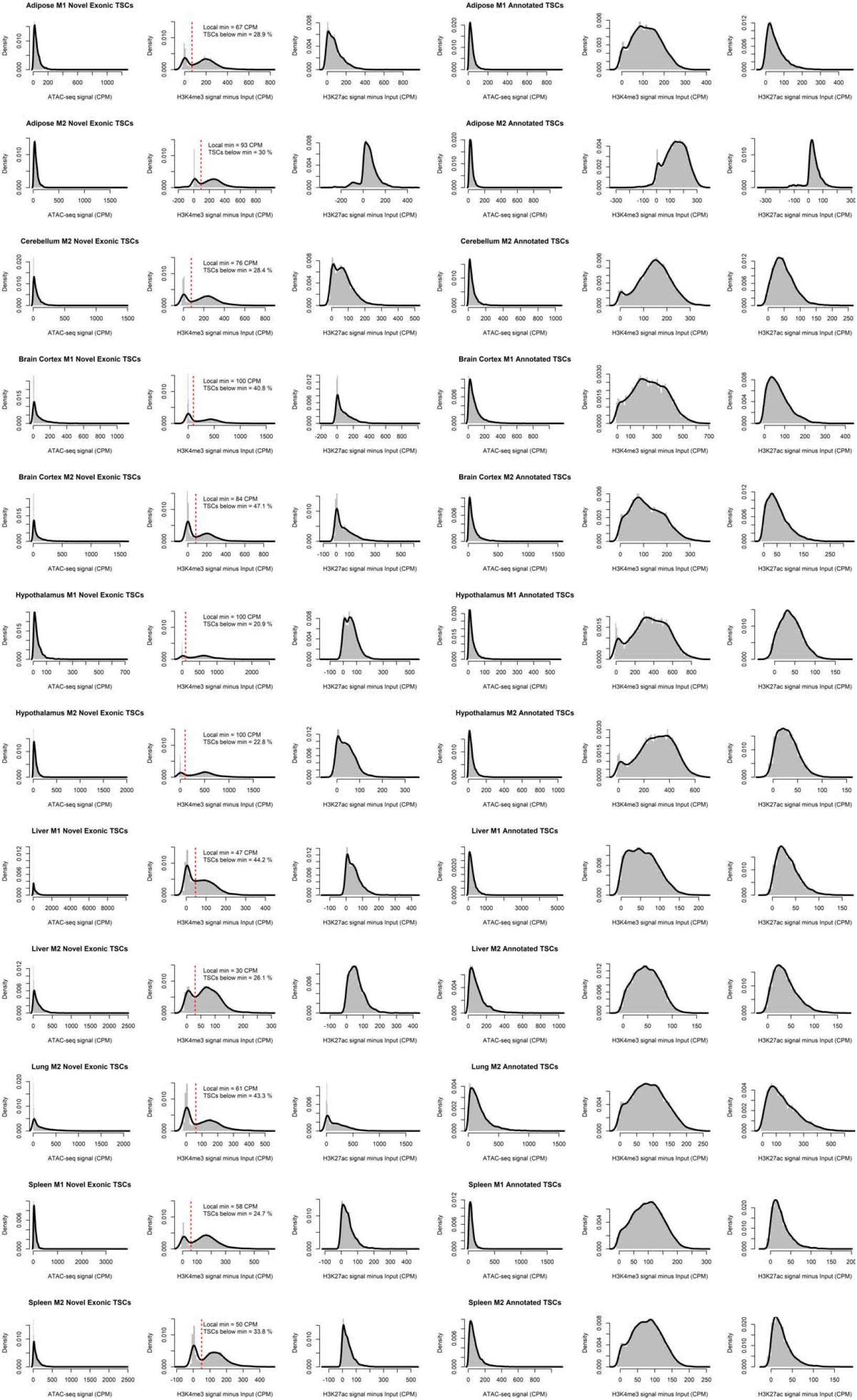
ATAC-seq, H3K4me3, and H3K27ac marks around novel TSCs in annotated exons (left) and known TSSs (right). The bimodal distribution of H3K4me3 signal allowed us to estimate that approximately 30% of the novel TSCs in annotated exons could be artifacts produced by incomplete cDNA synthesis or RNA degradation. TSCs under the local minimum of H3K4me3 signal in all samples were labeled in Supplemental Table S2.

### Unassigned TSCs

Many of the TSCs (n=44,125) showed no associations with any annotated lncRNA or protein coding genes, which could have surged as consequence of pervasive transcription (Wade and Grainger 2014). Pervasive transcripts are easily distinguished as they are non-coding, not limited by gene boundaries, and frequently antisense, so we proceeded to evaluate such aspects. Unassigned TSCs localized mainly to intergenic (58%) and intronic regions (38%), and were particularly abundant in the vicinity of genic TSCs, but on the opposite strand (Figure 4A), representing antisense RNA likely generated by bidirectional transcription (Engström et al. 2006). The search for antisense elements in regions 1.5 kb upstream and downstream of genic TSCs revealed 6,946 antisense TSCs associated with genic TSCs from 5,247 genes.

**Figure 4.**
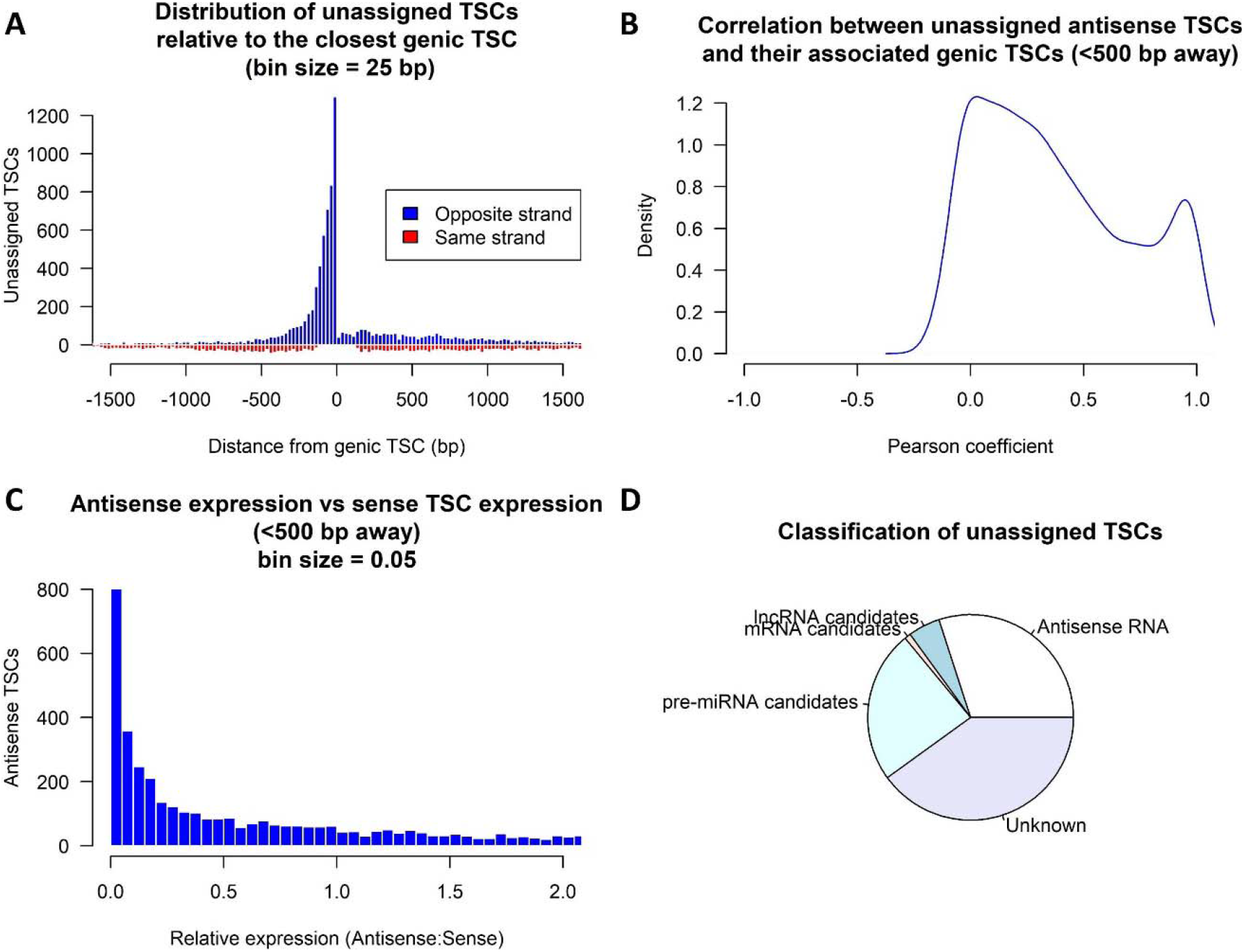
Identification of unassigned TSCs through bioinformatic approaches. (A) Distance between unassigned TSCs and their nearest genic TSCs. A high number of elements localized within 500 bp upstream of genic TSCs but on the opposite strand. (B) Correlation of expression from antisense TSCs (CPM > 3) with expression from sense TSCs. (C) Expression of the antisense variant (CPM > 3) relative to the sense variant. (D) Putative roles attributed to unassigned TSCs.

Interestingly, 60% of these cases occurred within 500 bp upstream of genic TSCs (Figure 4B), which is consistent with previous observations in mouse (58%) (Lepoivre et al. 2013). Some of these antisense TSCs were highly correlated with their sense complements, suggesting they were regulated by the same promoter (Figure 4C). In terms of expression, antisense TSCs generally contributed substantially fewer transcripts than sense TSCs (Figure 4D). In addition to antisense TSCs in proximity to sense TSCs, we identified 6,342 antisense TSCs in gene bodies, resulting in 13,942 antisense TSCs in total, and accounting for 30% of the unassigned TSCs.

To further characterize the remaining unassigned TSCs, we generated partial transcript models (Supplemental File S1) and predicted their coding potential based on *k*-mer frequencies, codon usage, and open reading frame coverage. As a result, we identified 2,306 TSCs associated with potential lncRNAs (Supplemental File S2) and 259 TSCs associated with potential mRNAs (Supplemental File S3). Lastly, we predicted pre-miRNA structures among the remaining models and identified 11,266 TSCs for candidate miRNA genes (Supplemental File S4), which is slightly over the estimated number of human miRNA candidates (Alles et al. 2019). After identifying TSCs for potential mRNAs, lncRNAs, and pre-miRNAs, the number of remaining unassigned elements decreased to 18,259. The epigenetic marks at these loci showed unclear patterns (Supplemental Fig S6), and the RAMPAGE signal failed to cluster samples by tissue except for a few cases (Supplemental Fig S7), suggesting the remaining TSCs represented technical artifacts. However, we could not discard the possibility that these TSCs corresponded to unannotated small RNAs, other types of RNAs, or repetitive elements.

**Supplemental Fig S6.**
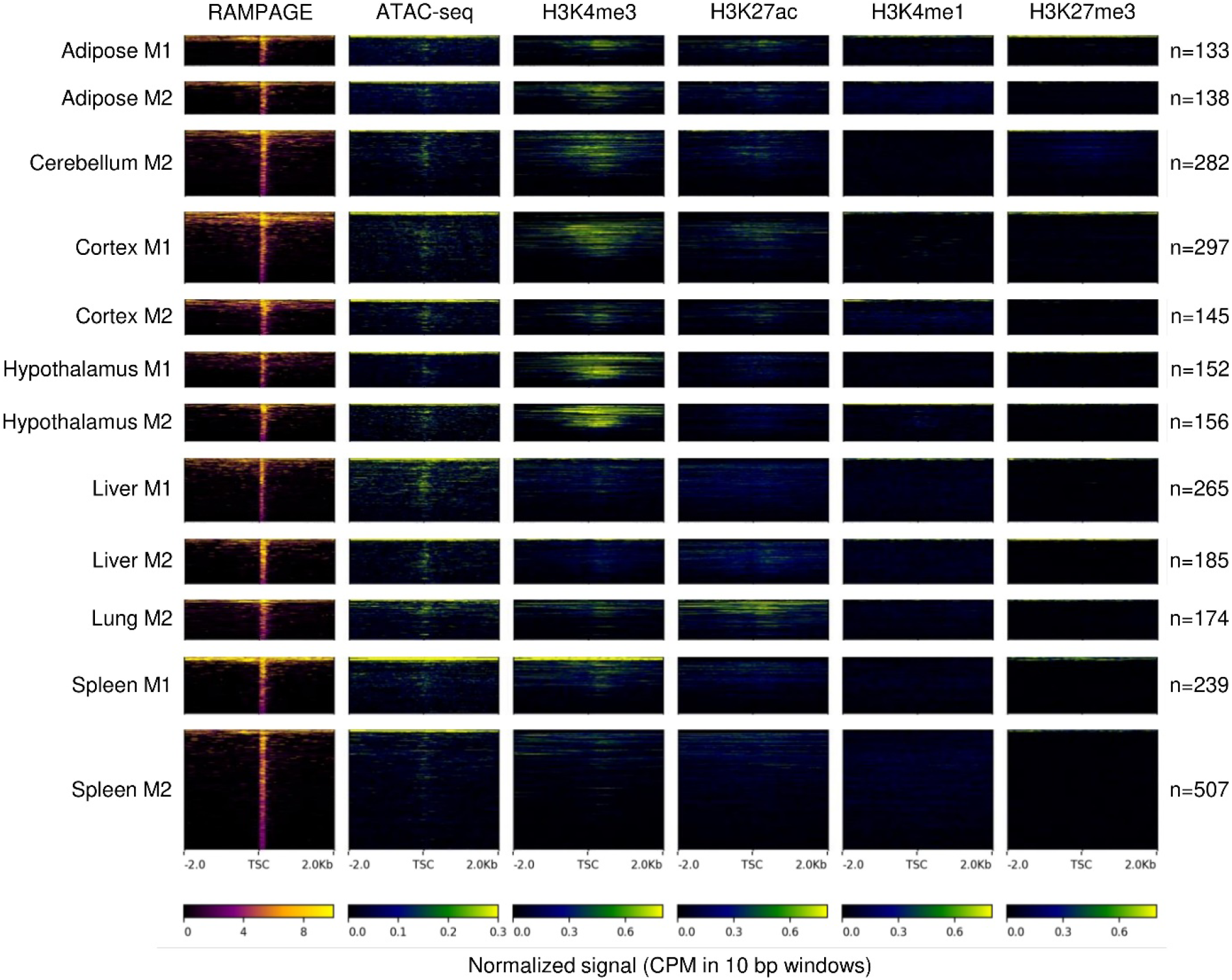
Epigenetic marks at the remaining unassigned TSCs after prediction of mRNAs, lncRNAs, and pre-miRNAs. From the unclear patterns, we concluded these sites comprised a heterogeneous population of RNAs with unclear functions and technical noise.

**Supplemental Fig S7.**
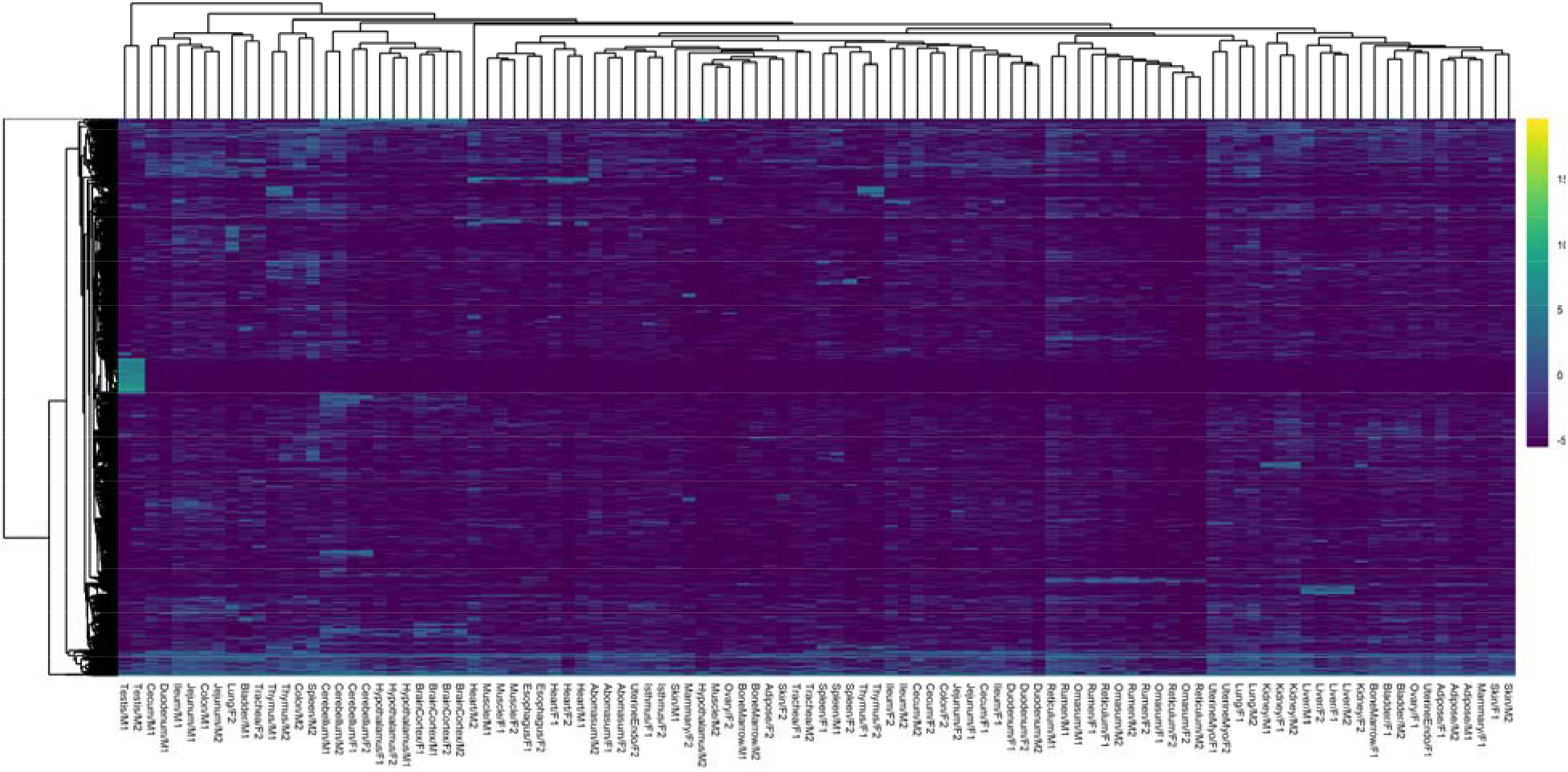
Hierarchical clustering based on RAMPAGE signal (variance-stabilized counts) at the remaining unassigned TSCs. Samples failed to cluster appropriately, suggesting many of these unassigned TSCs could be artifacts.

### Analysis of promoter activity and specificity

As RAMPAGE relies on the quantification of 5’ signal from mature transcripts, this technique also provides useful information for gene expression and promoter activity studies. To demonstrate this aspect, we first analyzed our samples by hierarchical clustering (Figure 5A). Samples grouped according to tissue and higher order structures, proving that the method was quantitative enough to separate samples appropriately. All the RAMPAGE information, including peak location, gene assignment, quality of the gene assignment, peak status by epigenetic signal and other features, as well as the number of tags in each sample, was implemented into a TSC atlas (Supplemental Table S2). Genic TSCs are also available as GTF file (Supplemental File S5). To evaluate the complexity of the dataset and to identify groups of potentially co-regulated elements, we generated a co-expression network based on the number of RAMPAGE tags in each sample (Figure 5B). The resulting network comprised 34 modules of co-expressed TSCs with different degrees of tissue-specificity, which varied from single tissues, such as trachea or thymus, to high-level structures, such as brain or digestive epithelium (Figure 5C, Supplemental Table S3). The largest module was highly correlated to testis and comprised 6,854 TSCs, followed by the brain module with 3,194 TSCs, and the intestines module with 1,888 TSCs. These numbers were consistent with studies in human and mouse, which have demonstrated that the testis expresses the largest number of tissue-specific genes (over twice as many as the second ranked tissue, brain cortex) (Djureinovic et al. 2014; Guo et al. 2005). Promoters from each module were significantly enriched in binding motifs for transcription factors associated with their respective tissues (Supplemental Fig S8). For instance, the Esophagus-Heart-Muscle module was enriched in binding motifs for all members of the myocyte enhancer factor-2 (*MEF2*) family, which are known to regulate muscle specific genes and muscle development (Gossett et al. 1989).

**Supplemental Table S2.** Promoter activity atlas. Location, gene assignment, assignment reliability, status by epigenetic signal, status by expression features, and expression (CPM) of each RAMPAGE peak in the dataset. Counts were normalized with respect to the total number of fragments associated to peaks in each sample. Assignment reliability was defined based on the maximum number of reads supporting a gene association: count ≥ 10 (high), 3 < count < 10 (medium), count ≤ 3 (low). The epigenetic status of each peak indicates whether peaks were under the local minimum of H3K4me3 signal used to estimate false positives by epigenetic signal, the status by assumptions column indicates whether the peak fits the criteria proposed to distinguish false positives (downstream location, high correlation to main TSC, and lower expression than main TSC).

**Supplemental Table S3.** Genic TSCs and their corresponding co-expression modules.

**Supplemental Fig S8.**
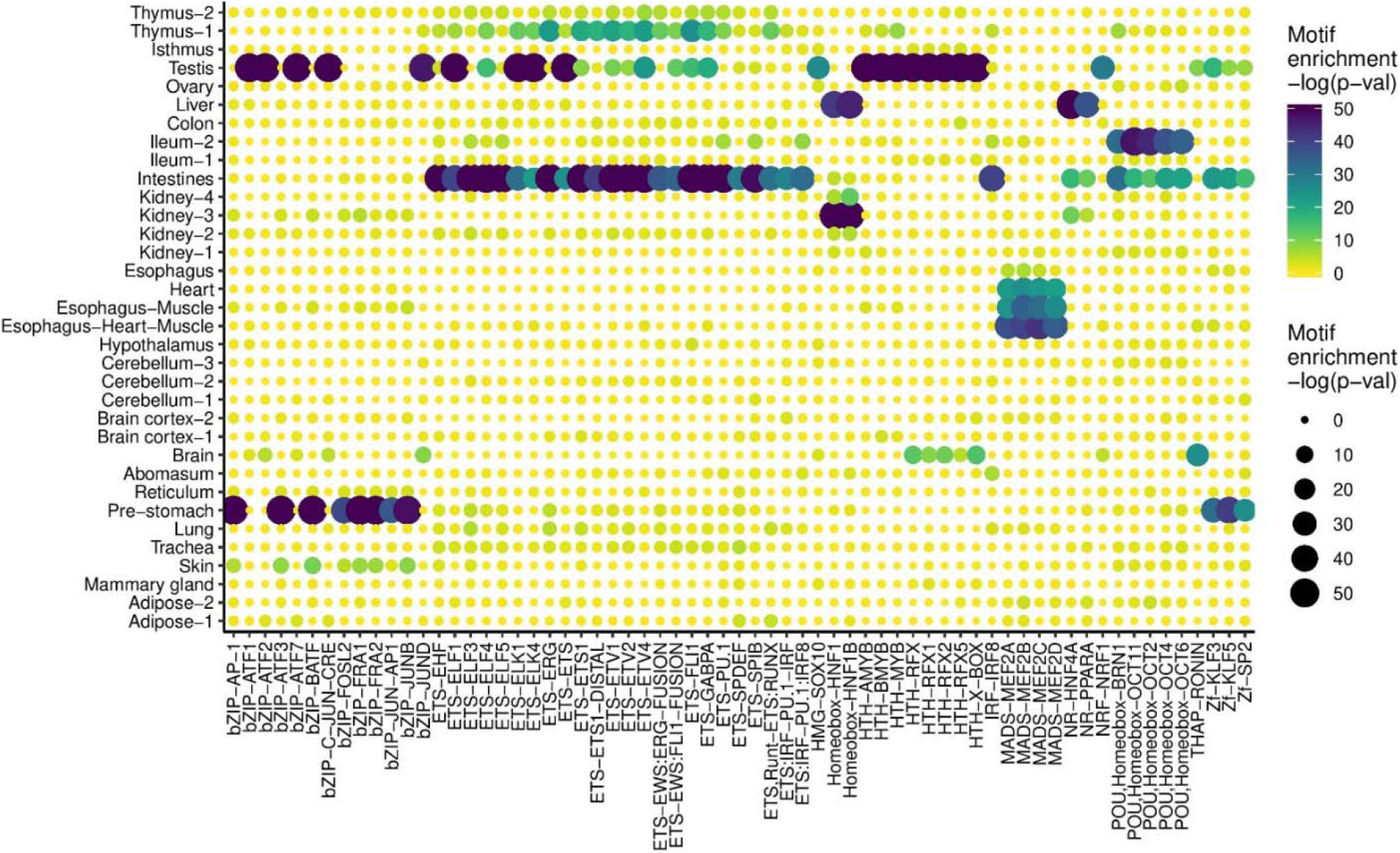
Binding motif enrichment in regions from 300 bp upstream to 100 bp downstream of co-expressed TSCs.

**Figure 5.**
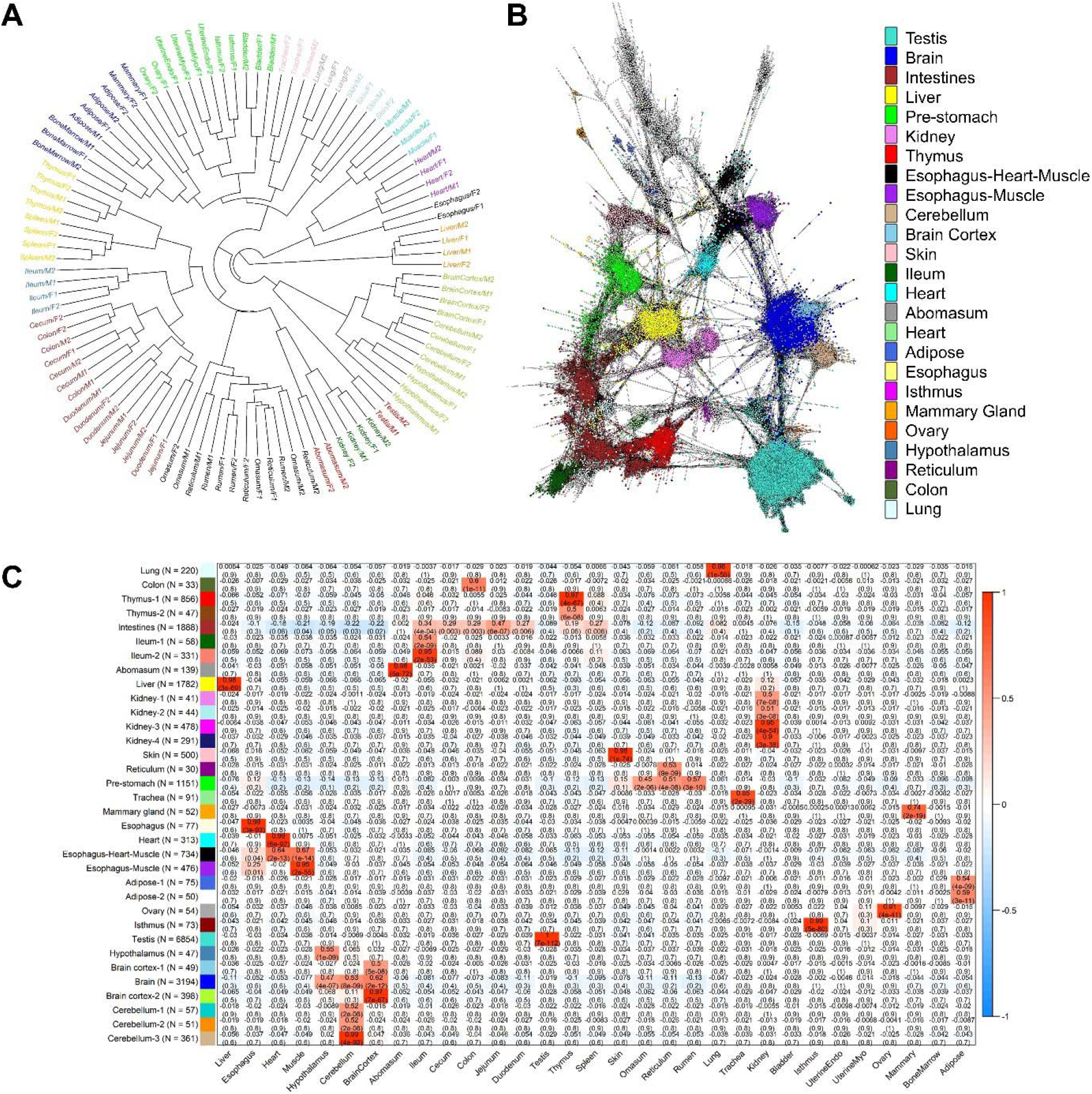
Promoter activity detected in 31 cattle tissues by RAMPAGE. (A) Sample dendrogram based on RAMPAGE signal. Samples grouped according to tissue, system and higher order structures. (B) TSC-to-TSC network generated based on Pearson correlations. This network demonstrates the diversity of tissue specific promoters in our dataset. Figure was generated using a minimum correlation of 0.75 in the Graphia v2.0 software (Freeman et al. 2020). (C) Modules of co-expressed TSCs indicating Pearson correlations to each tissue and p-values.

To validate the RAMPAGE signal from a quantitative perspective, we compared RAMPAGE counts to conventional RNA-seq gene counts in seven tissues from the same two male individuals. Estimates of gene expression by RAMPAGE were highly reproducible between biological replicates (average Pearson’s R = 0.94, SD = 0.03) (Supplemental Table S4, Supplemental Fig S9), consistent with the reproducibility of conventional RNA-seq (average Pearson’s R = 0.98, SD = 0.01) (Supplemental Fig S10). Absolute quantification of gene expression was comparable between the two techniques (average Pearson’s R = 0.76, SD = 0.03) (Supplemental Figs S11, S12), and detection of differentially expressed genes was strongly correlated between RNA-seq and RAMPAGE (average Pearson’s R = 0.9, SD = 0.05) (Supplemental Fig S13). Overall, these results suggest slight differences in global transcriptome measurement by RAMPAGE and RNA-seq, although both assays captured highly similar levels of differential gene expression.

**Supplemental Table S4.**
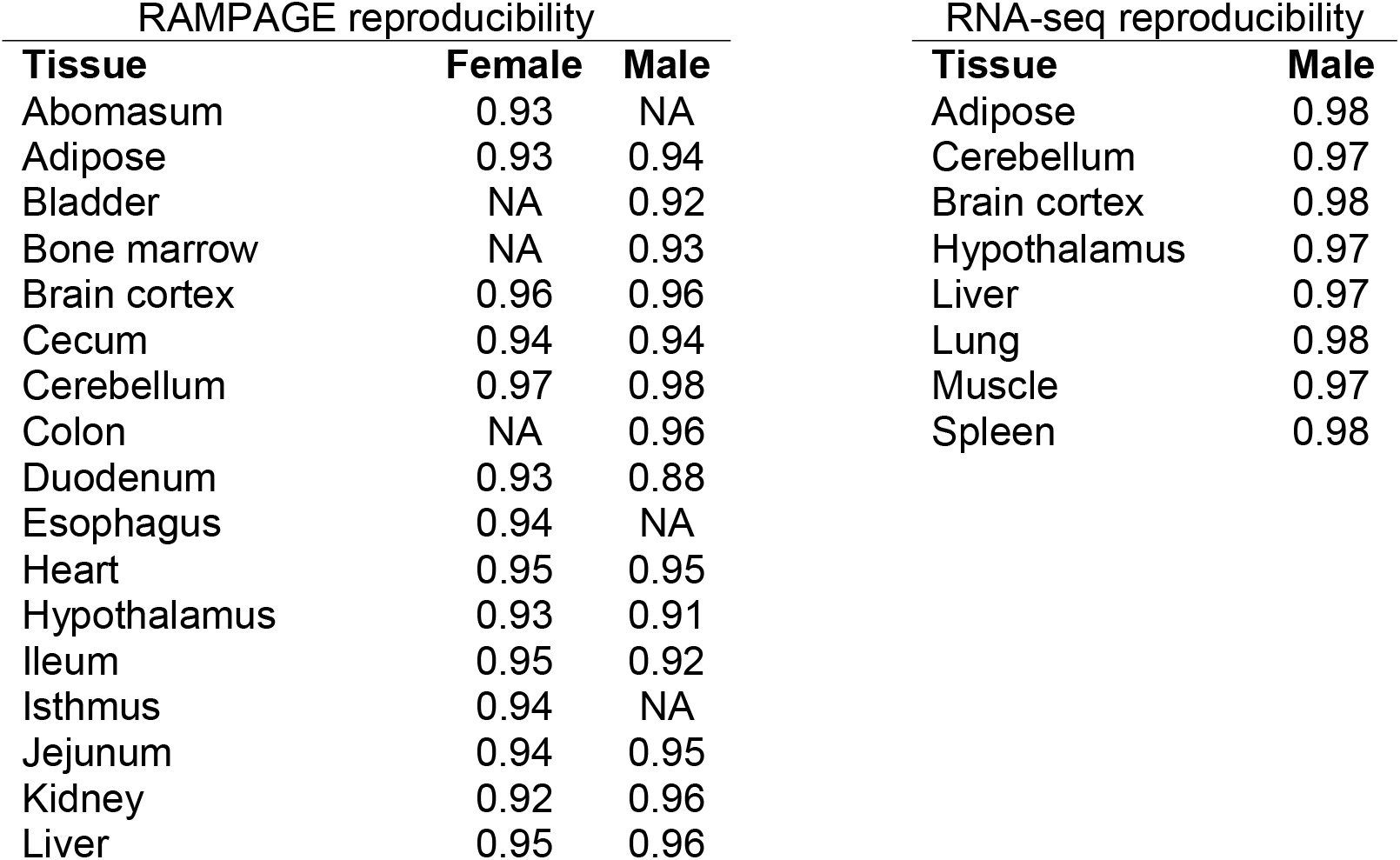
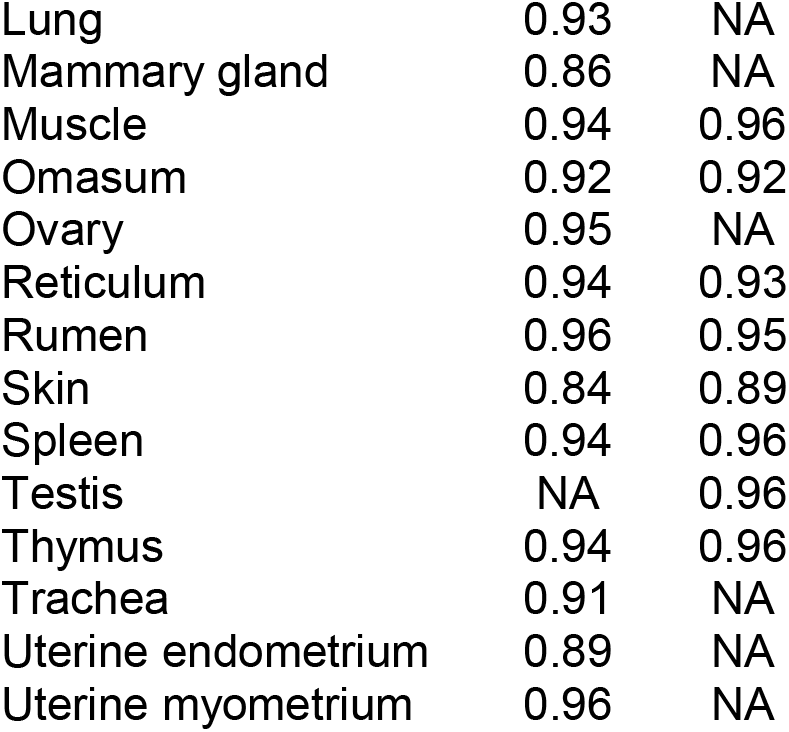
Reproducibility of gene expression between RAMPAGE and RNA-seq replicates. Based on Pearson correlations, estimates of gene expression by RAMPAGE were highly reproducible between biological replicates, consistent with reproducibility of conventional RNA-seq.

| RAMPAGE reproducibility |  |  | RNA-seq reproducibility |  |
| --- | --- | --- | --- | --- |
| Tissue | Female | Male | Tissue | Male |
| Abomasum | 0.93 | NA | Adipose | 0.98 |
| Adipose | 0.93 | 0.94 | Cerebellum | 0.97 |
| Bladder | NA | 0.92 | Brain cortex | 0.98 |
| Bone marrow | NA | 0.93 | Hypothalamus | 0.97 |
| Brain cortex | 0.96 | 0.96 | Liver | 0.97 |
| Cecum | 0.94 | 0.94 | Lung | 0.98 |
| Cerebellum | 0.97 | 0.98 | Muscle | 0.97 |
| Colon | NA | 0.96 | Spleen | 0.98 |
| Duodenum | 0.93 | 0.88 |  |  |
| Esophagus | 0.94 | NA |  |  |
| Heart | 0.95 | 0.95 |  |  |
| Hypothalamus | 0.93 | 0.91 |  |  |
| Ileum | 0.95 | 0.92 |  |  |
| Isthmus | 0.94 | NA |  |  |
| Jejunum | 0.94 | 0.95 |  |  |
| Kidney | 0.92 | 0.96 |  |  |
| Liver | 0.95 | 0.96 |  |  |

|  |  |  |
| --- | --- | --- |
| Lung | 0.93 | NA |
| Mammary gland | 0.86 | NA |
| Muscle | 0.94 | 0.96 |
| Omasum | 0.92 | 0.92 |
| Ovary | 0.95 | NA |
| Reticulum | 0.94 | 0.93 |
| Rumen | 0.96 | 0.95 |
| Skin | 0.84 | 0.89 |
| Spleen | 0.94 | 0.96 |
| Testis | NA | 0.96 |
| Thymus | 0.94 | 0.96 |
| Trachea | 0.91 | NA |
| Uterine endometrium | 0.89 | NA |
| Uterine myometrium | 0.96 | NA |

**Supplemental Fig S9.**
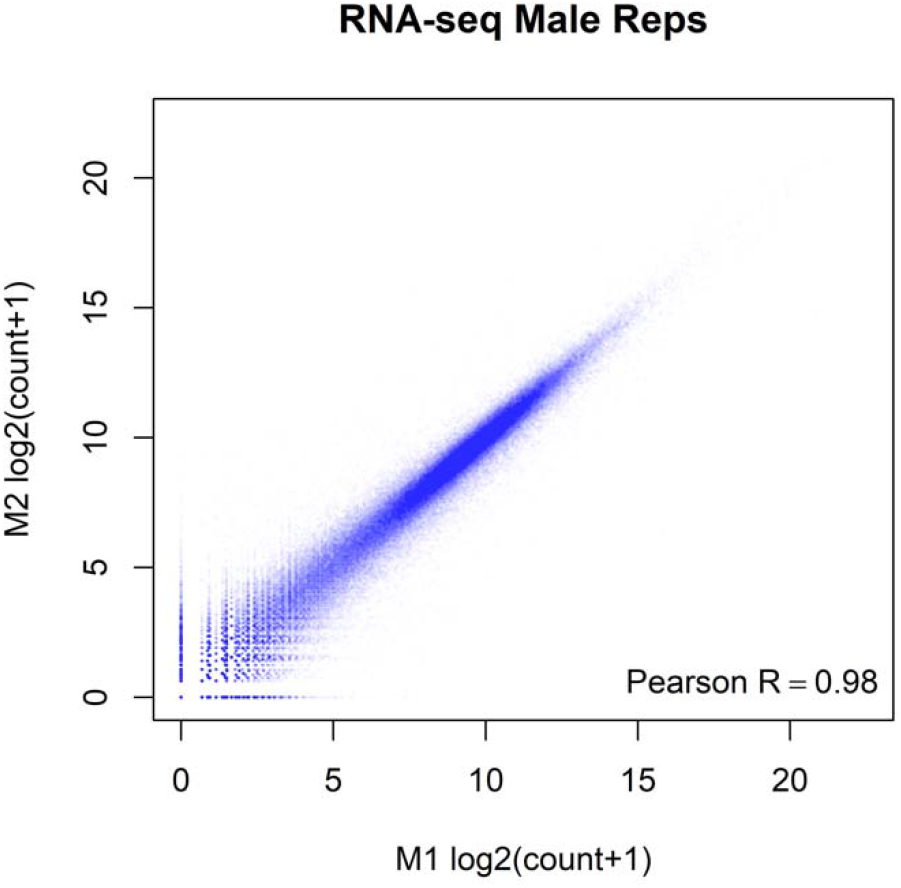
Reproducibility between RAMPAGE replicates. Counts were normalized by DESEQ2.

**Supplemental Fig S10.**
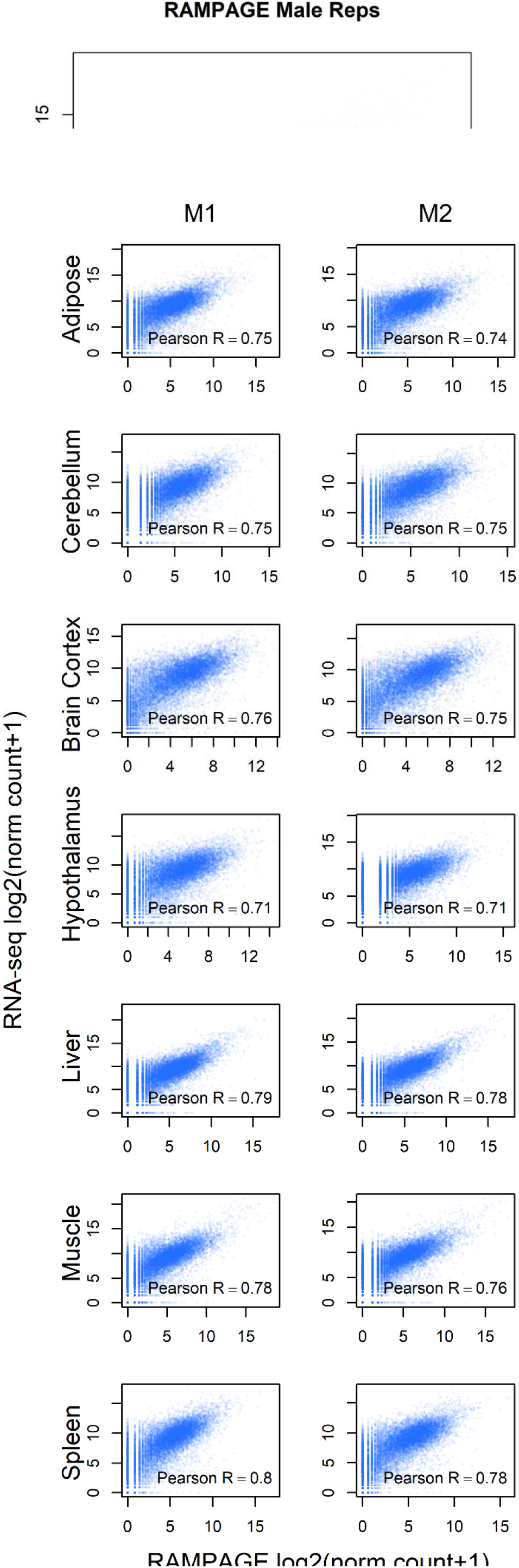
Reproducibility between replicates for RNA-seq counts. Counts were normalized by DESEQ2.

**Supplemental Fig S11.**
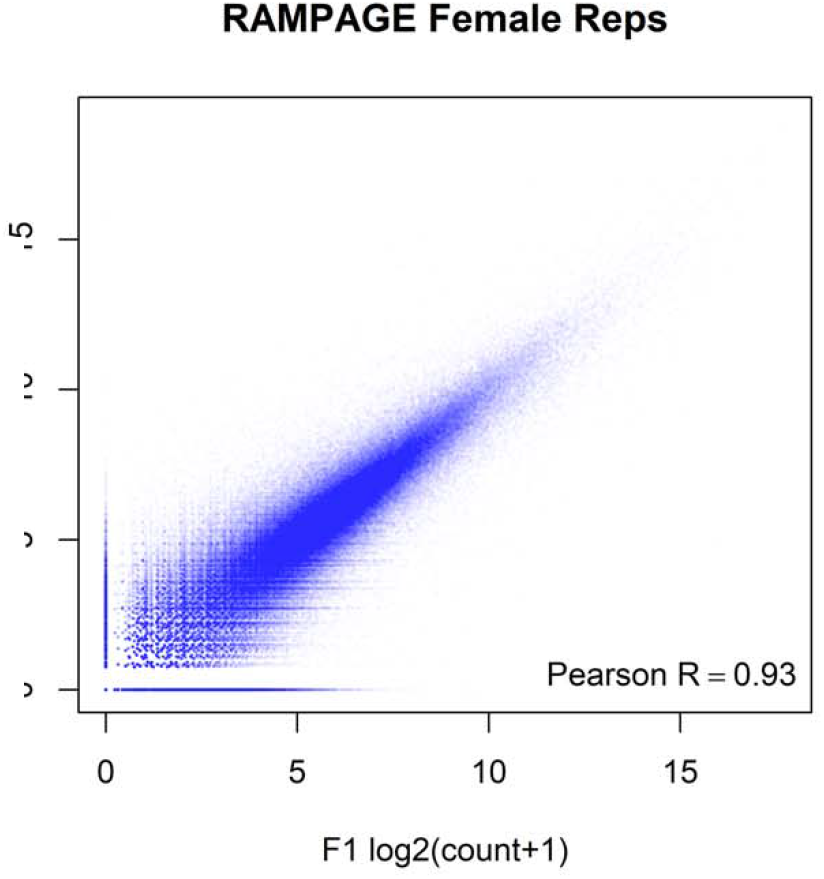
Pairwise correlations between RNA-seq and RAMPAGE log-transformed normalized counts. Moderately high correlations were consistently observed across samples. Counts were normalized by DESEQ2.

**Supplemental Fig S12.**
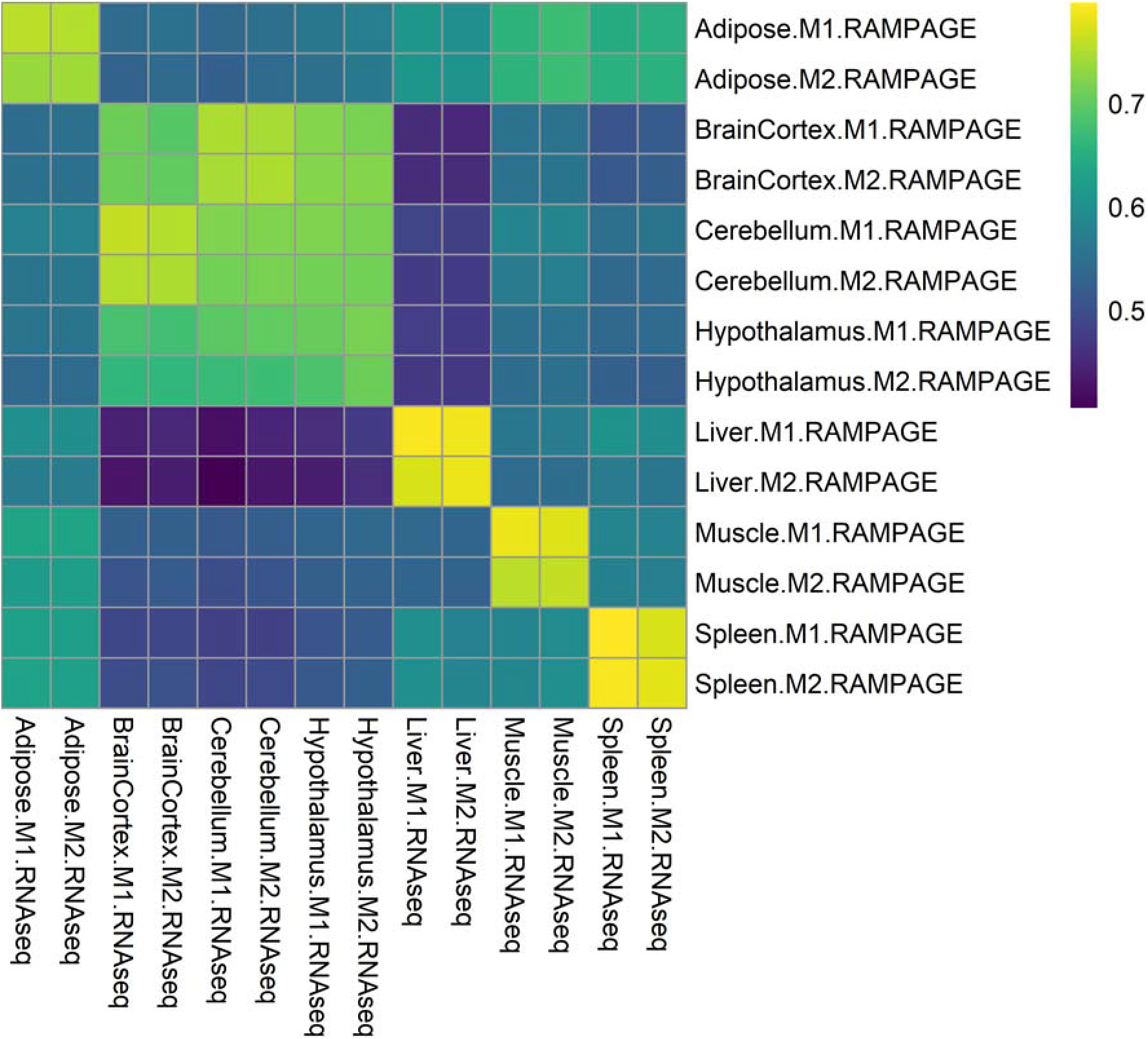
Pearson correlation between RNA-seq and RAMPAGE log-transformed normalized counts. High correlation was observed between samples from the same tissue.Counts were normalized by DESEQ2.

**Supplemental Fig S13.**
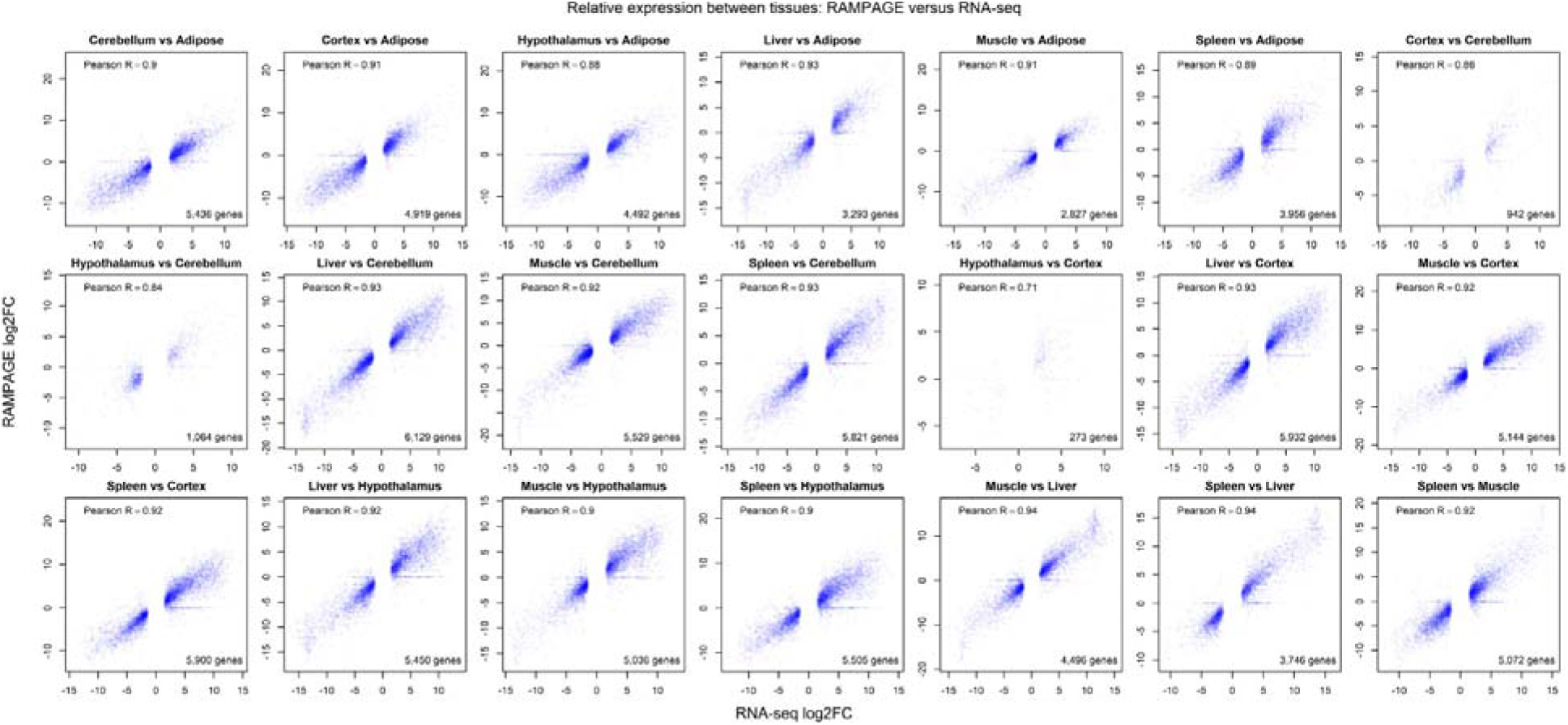
Pairwise correlations between logFC values obtained from RNA-seq and RAMPAGE gene counts. The analysis contemplates differentially expressed genes first identified by RNA-seq. Correlations were high except for brain tissues, where differences in gene expression were minimal.

After evaluating dataset complexity and validating the quantitative aspects of our data, we proceeded to evaluate the use of alternative promoters across tissues and the composition of each network module. Expression profiles from alternative TSCs of the same gene across the dataset were generally independent, as evidenced by the low correlation coefficients obtained in pairwise comparisons (Figure 6A). This observation suggested that most of the alternative TSCs were regulated by different biological mechanisms. Most of the TSCs were expressed in only one or a few tissues, with about 6,000 ubiquitous TSCs (Figure 6B). It is worth mentioning that some tissues, such as lung and trachea, expressed even more genes than testis and brain but showed few tissue specific TSCs in comparison (Figure 5C). To evaluate the expression of genes from tissue specific TSCs, we analyzed expression from 3,336 genes detected in at least two co-expression modules. Interestingly, we found 601 genes whose transcripts originated from alternative promoters in either testis or brain, which represented the highest use of alternative promoters between modules (Figure 6C, 6D, 6E). The Gene Ontology (GO) analysis of genes shared by testis and brain revealed a significant enrichment in terms related to cytoskeleton components, microtubule binding, ATP-binding kinase activity, synapse, cell-cell adhesion, microtubule-based movement and transport, and kinesin activity (Supplemental Table S5). We also found statistical enrichment in synapse pathways, including GABAergic synapse, cholinergic synapse, glutamatergic synapse, and serotonergic synapse, as well as critical pathways from the hypothalamus-pituitary-gonad (HPG) axis, such as GnRH and GABA signaling. The high number of members and use of alternative promoters found in these two tissues evidenced the complexity they acquired throughout evolution. Following the testis-brain combination, we found testis-intestines, with 357 genes, and testis-liver, with 325 genes using alternative promoters in each tissue. Genes shared by testis and intestines were highly enriched in GO terms related to cell cycle and division control, GTPase activity regulation, kinase activity, and additional groups of processes with lower enrichment (Supplemental Table S5). Genes shared by testis and liver, on the other hand, were slightly enriched in terms related to acyl-CoA-binding, mitochondrial components, and steroid and lipid metabolism (Supplemental Table S5).

**Figure 6.**
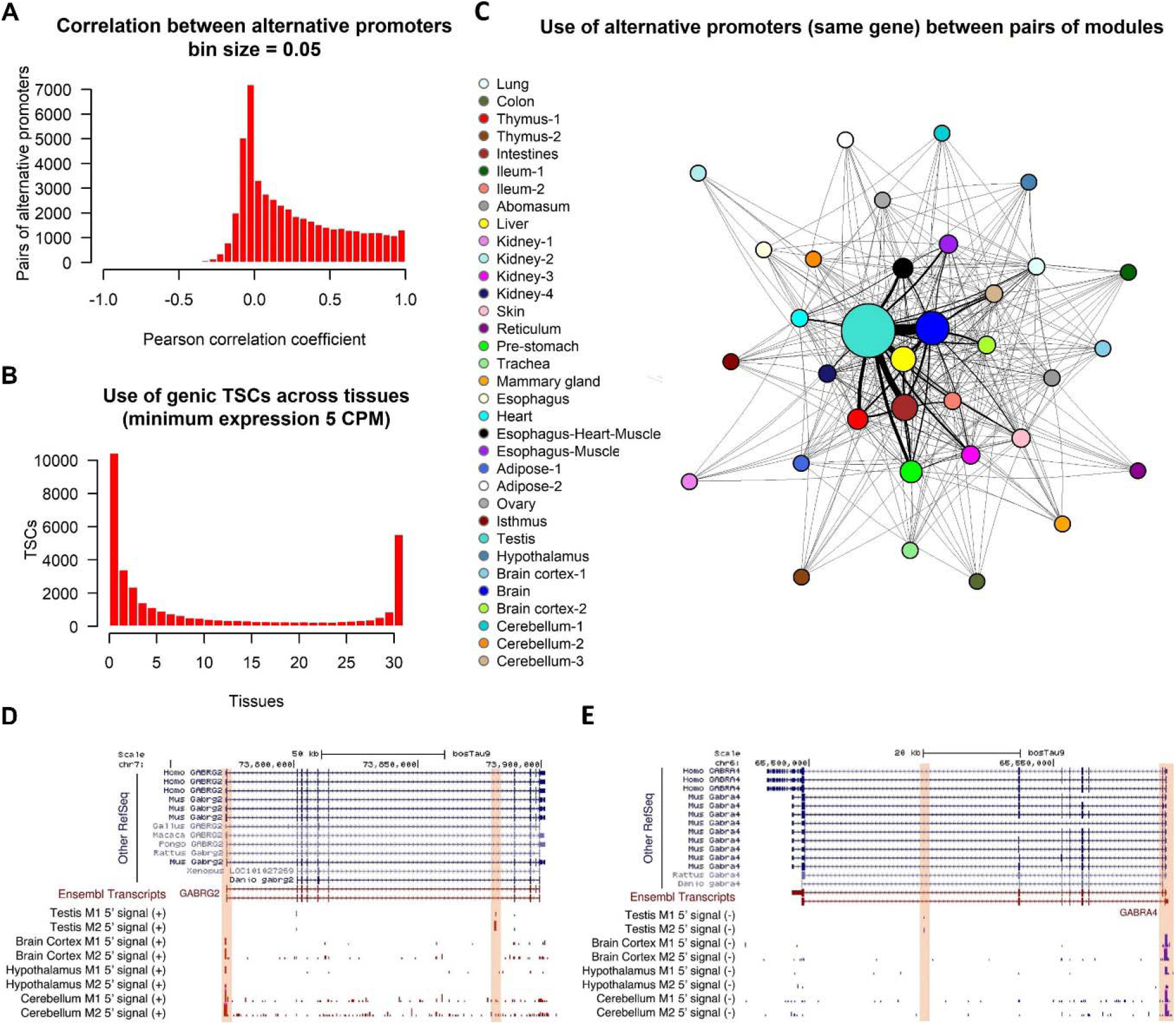
Usage of promoters across bovine tissues. (A) Correlation between pairs of alternative TSCs from the same gene (>5 CPM). Alternative TSCs were generally independent from each other. (B) Usage of TSCs across tissues. Most of the TSCs were expressed in only one or a few tissues, while about 6,000 TSCs were ubiquitously expressed. (C) Use of alternative TSCs between pairs of tissues. Sphere size represents the number of TSC members in the module, while edge thickness represents the number of common genes expressed from alternative (tissue specific) TSCs. (D, E) Examples of tissue specific promoters in brain and testis. The *GABRG2* and *GABRA1* genes are members of the GABAergic synapse pathway.

**Supplemental Table S5.** Functional annotation clustering of genes expressed by testis-brain, testis-intestines, and testis-liver from tissue specific TSCs. These tissue combinations showed the highest number of genes under such criterion.

## DISCUSSION

In this study, we analyzed a large collection of bovine tissues from both sexes using the RAMPAGE approach, identifying thousands of TSCs associated with protein coding and lncRNA genes. The diversity of analyzed tissues captured the expression of 18,312 previously annotated genes. However, no expression was detected for 5,016 genes, likely reflecting the absence of embryonic/fetal developmental genes in our collection of tissues from juvenile/adult animals. Thus, an additional analysis of samples at different stages of development should be performed to generate an even more comprehensive TSS annotation of the bovine genome. It is worth mentioning that samples from this study belonged to the same closed, inbred herd as the reference Line 1 cow. The genetic proximity between our samples and the reference genome likely contributed to the high mapping rates, while the low genetic variation provided by the inbred background of Line 1 likely contributed to the high correlations between samples.

Our dataset allowed us to identify several known and novel TSCs with a broad range of sizes. Size variation was expected as mammalian promoters are known to allow transcription to start from both well-defined positions (TATA box-enriched promoters) and broad intervals (CpG-rich promoters) (Carninci et al. 2006). In fact, we found that regions upstream of narrow TSCs were enriched with TATA-box elements, whereas regions upstream of broad TSCs were enriched in GC-box and CCAAT-box elements. In addition, novel TSCs were supported by experimental evidence that revealed the presence of chromatin accessibility and transcriptional activation marks (H3K4me3, H3K27ac) and near absence of H3K4me1, an epigenetic mark historically used for enhancer annotation (Heintzman et al. 2007), which implicated these novel TSCs as part of promoters. In addition to TSCs associated with annotated genes, we found many unassigned TSCs that supported the existence of pervasive transcription. A considerable number of unassigned TSCs were upstream of genic TSCs and seemed to be a consequence of bidirectional promoter activity. As shown in mouse (Lepoivre et al. 2013), transcripts from bidirectional promoters were more correlated to each other than transcripts from alternative TSCs of the same gene, suggesting a common regulatory mechanism. Furthermore, antisense transcripts presented low expression levels in comparison to their complementary sense transcripts, supporting the concept of pervasive transcription by erratic behavior of the polymerase.

Even though the RAMPAGE method has important advantages over other 5’-seq techniques for TSC identification (Adiconis et al. 2018), including CAGE (Takahashi et al. 2012) due to extensive use of paired-end information, false TSCs were expected to arise as products of incomplete cDNA synthesis and mRNA degradation. We estimated that about 30% of the 13,209 novel TSCs in annotated exons could be false positives as shown by the low H3K4me3 signal. These elements could have been caused by re-capping, which has been reported in RNAs with truncated 5’-ends (Otsuka et al. 2009) and within the body of mRNAs (Affymetrix/Cold Spring Harbor Laboratory ENCODE Transcriptome Project 2009). However, due to the low number of tissues with epigenetic information in comparison to the RAMPAGE dataset, our evidence was insufficient to identify and discard all potential artifacts. A global estimation based on the assumptions that artifacts are most frequent in exons (downstream of the strongest TSC), expressed to a lower extent (< 25%), and correlated to the strongest TSC for being part of the same transcript (Pearson > 0.5), suggested that 4,059 (8.7%) of the genic TSCs could be false. This estimation supported our initial estimation by epigenetic signal in the reduced validation dataset. However, the identification of these sites needs to be corroborated by additional epigenetic information from other tissues.

The analysis of co-expression networks revealed a high number of genes expressed in both brain and testicle under different tissue-specific TSCs. The high similarity between brain and testis expression profiles was reported years ago in mouse and human (Guo et al. 2005, 2003). The mechanisms and epigenetic factors contributing to the projection of the brain expression program on the testis are unclear, although intriguing. It has been proposed that the hypothalamus-pituitary-gonad (HPG) could be implicated in the maintenance of this similarity, given the importance of the brain expression program for human speciation and transmission to the offspring (Guo et al. 2005). Our study, however, demonstrated that although brain and testis share a high number of genes, transcripts are expressed from tissue-specific promoters. These promoters could be generating isoforms with different and unique functions. This information could be valuable to understand the regulatory mechanisms operating behind tissue-specific expression profiles.

With our set of new TSCs, the bovine ratio of TSS to genes reached a level comparable to the human and mouse genomes. Hopefully, this annotation will improve our understanding of the genetic control of complex livestock traits, complementing studies from other members of the FAANG community. In addition, our data will improve the connection between genotypes and phenotypes to exploit predictive models in the breeding field; provide new biological explanations for genetic diseases reportedly unexplained based on other annotations; and contribute to the study of genome evolution through comparative studies with other species.

## METHODS

### Sample collection

Samples from abomasum, adipose, bladder, brain cortex, bone marrow, cecum, cerebellum, colon, duodenum, esophagus, heart, hypothalamus, ileum, isthmus, jejunum, kidney, liver, lung, skeletal muscle, mammary gland, omasum, ovary, reticulum, rumen, skin, spleen, testis, thymus, trachea, uterine endometrium and uterine myometrium were collected from two female and two male 14-month old Line 1 Herefords provided by Fort Keogh Livestock and Range Research Lab. Samples belonged to the same closed, inbred herd as the original Line 1 reference cow.

Sample collection was carried out with all the necessary permissions, following Protocol for Animal Care and Use #18464, approved by the Institutional Animal Care and Use Committee (IACUC), University of California, Davis. Cattle were slaughtered by captive bolt under USDA inspection at the University of California, Davis, and samples were collected within one hour post euthanasia. Collected tissue samples were flash frozen in liquid nitrogen and stored at −80 °C until processing.

### RAMPAGE library preparation

Frozen tissues kept at −80°C were homogenized with a mortar and pestle in liquid nitrogen. Total RNA was extracted using TRIzol (Invitrogen, Carlsbad, CA) followed by a column clean-up using the Direct-zol RNA Mini Prep Plus kit (Zymo Research, Irvine, CA) and performing an in-column DNA digestion. Integrity of the DNase-treated RNA was verified on the Experion electrophoresis system (Bio-Rad, Hercules, CA). The RNA quality indicator (RQI) varied from 7.5 to 9.7, indicating good quality RNA, excepting Liver from Male A which had a borderline RQI of 6.9.

Following the method described by Batut & Gingeras, (2013), DNase-treated RNA was subjected to terminator digestion (TEX enzyme, Epicentre) to degrade 5’-monophosphate RNA (16S and 18S ribosomal RNA). Random primers bearing an adapter sequence overhang were annealed to the RNA, which was then subjected to reverse transcription, leaving a string of terminal cytosines. Template–switching oligonucleotides (TSO) containing the library identification barcodes were then hybridized to these terminal cytosines, prompting the reverse transcriptase to switch templates and add the TSO sequence to the end of the newly synthesized cDNA. The TSO bears an Illumina adaptor sequence, such that only 5’-complete cDNAs are amplifiable, whereas non-5’ complete molecules are not. Libraries were then quantified by qPCR to pool multiple samples for sequencing.

Libraries were pooled in equimolar amounts and 5’ caps were oxidized with sodium periodate. To prepare for cap trapping, 5’caps were biotinylated, and RNA molecules were pulled down using streptavidin magnetic beads. Finally, pooled libraries were PCR-amplified and size-selected using AmPure XP magnetic beads. Final pooled libraries were quantified in the Nanodrop spectrophotometer (Thermo Fisher Scientific, Wilmington, DE, USA) and the Bioanalyzer 2100 High-Sensitivity DNA chip (Agilent Technologies, Santa Clara, CA, USA). Libraries had the expected size range of 300 to 1000 bp, and were sequenced on an Illumina HS 4000 system, generating paired end reads of either 100 or 150 bp.

### RNA-seq data analysis

Data for RNA-seq (Kern et al. 2018) were downloaded from Gene Expression Omnibus (accession number GSE158430). Reads were trimmed using Trimmomatic (v0.33) to remove Illumina sequencing adapters and low-quality bases. Settings for adapter removal included a maximum of two mismatches, a palindrome clip threshold of 30, and a simple clip threshold of 10. Leading and trailing bases of quality less than 3 were removed. Sliding window trimming was conducted using a window size of 4 and minimum average quality of 15. After processing, reads shorter than 36 bp were discarded. Samples were aligned to the ARS-UCD1.2 *Bos taurus* reference genome using STAR v2.6.1 (Dobin et al. 2013) with options -- outFilterScoreMinOverLread 0.85 and --seedSearchStartLmax 30.

### RAMPAGE data analysis

Samples were aligned to the ARS-UCD1.2 *Bos taurus* reference genome using STAR in two-pass mode. The first six nucleotides of the first read (barcodes) were removed prior to alignment as part of the demultiplexing step. The last 15 nucleotides of the second read corresponded to the RT-primer, thus they were soft-clipped using the --clip5pNbases option during the alignment step. Only uniquely mapped reads were kept for subsequent analyses. In order to improve the specificity of the peak calling and the accuracy of transcript quantification, duplicates were marked and removed based on the alignment coordinates and the last 15 nucleotides of the second read (random primer), which were used as a pseudo-random single-molecule barcode to distinguish true duplicates. Alignments from the same sample were merged prior to peak calls. A preliminary assessment of data quality was carried out using hierarchical clustering of gene counts generated by featureCounts v1.6.2 (Liao et al. 2014) prior to peak calling. Samples showing anomalous expression profiles compared to other biological replicates from the same tissue were discarded from subsequent analyses.

### RAMPAGE peak calling

The 5’ coordinate of each cDNA was recorded and compiled into a single wiggle file as a quantification of every possible TSS in the dataset. Then, the same was done for every genomic position covered by downstream reads to create a signal background for the peak calling algorithm. Peak calling was conducted using the Python script developed by Batut et al. (2013) with a dispersion parameter of 4, a background weight of 0.6, and an FDR of 1E-8 in the combined dataset (Supplemental Code 1). Briefly, the algorithm works as a sliding window that, for each position in the genome, assesses the statistical enrichment of 5’ signal against a negative binomial background distribution. The coverage by downstream reads in such window is used to subtract a pseudo-count from the 5’ signal so that significance is harder to achieve at highly transcribed exon positions. Then, neighboring windows are merged and trimmed at the edges down to the first base with signal.

To increase the sensitivity for tissue specific peaks, each tissue was also analyzed separately using a dispersion parameter of 0.2, a background weight of 0.8, and an FDR of 1E-8. Tissue-specific TSCs undetected in the combined dataset, supported by at least 3 independent RAMPAGE tags, located outside exons, and conserved in at least two biological replicates, were incorporated into the global list of TSCs.

### Gene assignment

TSCs were assigned to genes using featureCounts in stranded mode (-s 1) and the Ensembl v95 annotation. Gene associations supported by less than three independent fragments were considered weak associations. To exclude spurious associations generated by run-off transcription, i.e. transcription that failed to terminate at the right site and continued over a downstream gene, associations to alternative genes supported by 5-fold fewer reads than the most supported gene were disregarded. For quantification purposes, the RAMPAGE signal was normalized by library size (total number of reads associated with TSCs). Genes were considered as expressed when their expression (normalized with respect to all the RAMPAGE tags assigned to genes) reached a minimum of 3 counts per million (CPM).

### Expression quantification by RNA-seq and RAMPAGE

For comparison between RAMPAGE and RNA-seq, RAMPAGE gene expression was quantified using the number of 5’ tags in TSCs attributed to each gene. Genes with less than 10 total counts across all samples were removed from the analysis prior to normalization by DESeq2 (Love et al. 2014) to eliminate background noise. Normalized counts were log-transformed and used to calculate Pearson correlations between biological replicates for each tissue. Log-transformed normalized counts from all tissues were then pooled to calculate the overall Pearson correlations between male replicates in all tissues and female replicates in all tissues. Gene expression was evaluated from RNA-seq data for a subset of seven tissues. Raw expression values for genes in the *Bos taurus* annotation (Ensembl v95) were calculated from uniquely mapped reads using the SummarizeOverlaps function from the GenomicAlignments R package (Lawrence et al. 2013) with the ‘union’, paired-end, and stranded options. Genes with less than 10 total counts across all samples were removed from the analysis prior to normalization to eliminate background noise. Gene counts were then normalized by DESeq2 and log-transformed. Log-transformed normalized counts from all seven tissues were pooled to calculate the overall Pearson correlation between male replicates. For the seven tissues for which RNA-seq data were available, log-transformed normalized counts were used to calculate Pearson correlations between RNA-seq and RAMPAGE. Finally, differential gene expression was compared between RNA-seq and RAMPAGE. From RNA-seq counts, differentially expressed genes (DEG) were identified for each pair of tissues using DESeq2 and lfcShrink (type=”apeglm”, lfcThreshold=1), given an s-value < 0.005. The log fold change (logFC) of these DEG were then calculated by DESeq2 according to RNA-seq and RAMPAGE signal and used to calculate Pearson correlations.

### ATAC-seq and ChIP-seq data analysis

ChIP-seq and ATAC-seq data (Kern et al. 2018) were downloaded from Gene Expression Omnibus (accession number GSE158430). ATAC-seq reads were trimmed with trim_galore! v0.4.0 (github.com/FelixKrueger/TrimGalore) (-q 20 -a CTGTCTCTTATA -stringency 1 –length 10) and aligned to the ARS-UCD1.2 genome with BWA-MEM (Li and Durbin 2009). Duplicate alignments were removed using Picard Tools v2.9.1 (broadinstitute.github.io/picard). Low-quality alignments (q < 15) were removed with SAMtools v1.9 (Li et al. 2009). ChIP-seq reads were processed following the original study (Kern et al. 2018). To visualize signal, the bamCoverage function of the deepTools v3.2.0 software (Ramírez et al. 2016) was used to calculate normalized coverage (CPM) in 10 bp windows genome-wide. For ChIP-seq, signal from input libraries was first subtracted with deepTools bigwigCompare. Normalized signal from ATAC-seq and ChIP-seq libraries was plotted alongside RAMPAGE 5’ signal on the UCSC genome browser. The deepTools computeMatrix and plotHeatmap functions were also used to plot normalized epigenetic signal at TSCs, plotting the median signal above heatmaps and using the interpolation method “bilinear” for smoothing. The average epigenetic signal (CPM) at TSC was calculated with the bigWigAverageOverBed function from the UCSC Genome Browser utilities.

### Identification of binding motifs

The abundance of TATA, GC and CCAAT boxes in regions 200 bp upstream to 200 bp downstream of novel TSCs was visualized using the seqPattern R package (https://bioconductor.org/packages/release/bioc/html/seqPattern.html. The search was carried out using the TATA-binding protein (Tbp) model included in the package, the MA0079.2.pfm JASPAR model for specificity protein 1 (Sp1) and the MA0060.3.pfm JASPAR model for nuclear transcription factor Y (Nfy).

Statistical enrichment was evaluated using the ‘findMotifsGenome.pl’ tool from HOMER v4.10.4 (Heinz et al. 2010). Background consisted of either random sequences from the rest of the genome or sequences from the rest of the gene body depending on the location of the TSC evaluated (promoters and intergenic regions, or exons and introns, respectively). The search was conducted on sequences 300 bp upstream to 100 bp downstream of the TSCs using binding motifs for Tbp, Sp1 and Nfy available in the HOMER database.

### Attribution of unassigned TSCs

Unguided partial transcript models were generated for RNAs derived from unassigned TSCs using StringTie v2.0.4 (Pertea et al. 2015) in --fr stranded mode, a minimum isoform fraction of 0, a minimum assembled transcript length of 30, a minimum of one read per bp coverage, and trimming based on coverage disabled. Antisense RNAs mapping to the gene body were identified using featureCounts in the opposite stranded mode (-s 2). Putative protein coding and lncRNA genes were predicted based on *k*-mer frequency, codon use, and ORF coverage using the FEELnc software (Wucher et al. 2017). Lastly, prediction of pre-miRNA candidates was carried out using the miRNAfold tool (Tav et al. 2016) with a sliding window size of 60 bp and FASTA files containing the remaining partial transcripts as input.

### Evaluation of alternative promoter usage

Correlations between alternative TSCs from the same gene were evaluated considering highly expressed genic TSCs (>5 CPM normalized with respect to the number of RAMPAGE tags assigned to genic TSCs) through the Pearson coefficient.

The use of alternative promoters was analyzed through signed co-expression networks generated by the WGCNA R package (Langfelder and Horvath 2008). To reduce noise, networks were constructed for TSCs with a minimum expression of 3 CPM in at least two samples of any tissue. The adjacency matrix was generated using variance-stabilized counts as input, biweight mid-correlations and a soft-thresholding power of 12. Modules were determined using the Dynamic Hybrid Tree Cut algorithm with a sensitivity (deepSplit) of 2 and PAM mode activated.

### Gene ontology analysis

Functional annotation clustering was conducted using official gene symbols in DAVID bioinformatics v6.8 (Huang et al. 2009). We used the *Homo sapiens* model as background due to its more comprehensive pathway annotation. The analysis was carried out using the default categories (OMIM_DISEASE, COG_ONTOLOGY, UP_KEYWORDS, UP_SEQ_FEATURE, GOTERM_BP_DIRECT, GOTERM_CC_DIRECT, GOTERM_MF_DIRECT, BBID, BIOCARTA, KEGG_PATHWAY, INTERPRO, PIR_SUPERFAMILY, SMART) and medium stringency.

## DATA ACCESS

All raw sequencing data generated in this study have been submitted to the NCBI BioProject database (https://www.ncbi.nlm.nih.gov/bioproject/) under accession number PRJNA630504. As samples were submitted using their FAANG nomenclature, sample M1 is found as M08, sample M2 is found as M22, sample F1 is found as F05, and sample F2 is found as F12. The Python script used to identify RAMPAGE peaks is available as supplemental material (Supplemental Code S1). Additionally, we generated a GTF file with partial transcripts (Supplemental File S1), a BED file with TSCs associated with lncRNAs (Supplemental File S2), a BED file with TSCs associated with unannotated genes (Supplemental File S3), a BED file with TSCs associated with pre-miRNAs (Supplemental File S4), and a GTF file with genic TSC annotations (Supplemental File S5).

## AKNOWLEDGEMENTS

Funding for this project was provided by USDA-NIFA-AFRI grant no. 2017-67015-26297 to PJR. ChIP-seq data was generated as part of USDA-NIFA-AFRI grant no. 2015-43567015-22940 to HZ and PJR. MMH was supported by a USDA-NIFA National Needs Fellowship Grant (USDA-NIFA Competitive Grant Project no. 2014-38420-21796) and an Austin Eugene Lyons Fellowship.

