## Supplemental_Code_S1 for "Transcription initiation mapping in 31 bovine tissues reveals complex promoter activity, pervasive transcription, and tissue-specific promoter usage": Supplemental_Code_S1.py

```
# rampagePeakCaller.py
# Philippe BATUT
# Gingeras Lab, CSHL 
# Sept. 22, 2011


# This script takes as input a WIGGLE file describing the density of RAMPAGE read 5' ends 
# over the genome. For a sliding window of size k sliding through each position across the 
# genome, it computes the probability of obtaining the observed number of RAMPAGE tags under 
# the null model. The null model is a negative binomial distribution whose mean is the 
# observed average density of tags per window, and whose dispersion parameter is user-defined. 
# 
# Optionally, it can also be provided with a second wiggle file describing the density of 
# RAMPAGE downstream reads (i.e., the read of the mate pair that starts at the RT priming 
# site). The downstream read coverage is used to subtract a pseudocount attributable to 
# background signal prior to computing the p-value for each window. The weight to be given 
# to this backgroung estimate is a user-defined parameter. 
# 
# The p-values are corrected by the method of Benjamini & Hochberg and thresholded to allow 
# for an FDR below a value defined by the user. All significant windows within a distance d 
# of each other are then merged, which generates the final peaks. 
# 
# 
# INPUT:    - Argument 1:		File providing the size of each chromosome in the reference 
#								genome (Format <chrName\tchrSize\n>). 
#			- Argument 2: 		RAMPAGE tag density file (Wiggle). 
#								The track line for each strand is required to contain a name
#								field ending with "_+" or "_-", from which the strand is 
#								deduced. 
#
#
# OPTIONS:  --width (-w): 			Sliding window width. Default: 15 bases. 
#			--NBinom (-n): 			Dispersion parameter of the negative binomial. Default: 0.6. 
#           --read2-background:		Downstream read density file (wiggle). 
#           --bg-weight (-w)		Weight to be given to the background signal. Default: 0.5. 
#           --fdr (-f):				False discovery rate. Default: 10^(-6). 
#           --exclude (-x):			Coma-separated list of chromosomes to exclude. Default: "chrM". 
#			--distance (-d): 		Maximum distance for merging neighboring windows into a single peak.
# 			--threads (-t): 		Number of threads to be used. Default: 1. 
#
#
# In our experience, the dispersion parameter (--NBinom) and the FDR (--fdr) are the most critical
# ones to change when optimizing the specificity of the peakcalling.  


 
from optparse import OptionParser
parser = OptionParser()
import os 
import sys 
import time 
import math 
import numpy 
import scipy.stats
import multiprocessing   
start_time = time.time()


###################################################################################################################
###################################################################################################################


######################## DEFINE FUNCTION TO LOAD & PARSE CAGE DATA FROM WIGGLE FILE:

def load_rampage_data(rampage_file_name, excluded_chr):
	rampage_file = open(rampage_file_name)
	parsed_rampage_data = {} 
	chr_tag_counts = {} 
	## Load RAMPAGE signal data:
	for line in rampage_file:
		if line.startswith('track'):
			trackLine = line.rstrip().split(' ') 
			strand = None 
			for field in trackLine: 
				if field.startswith('name='): strand = field.split('_')[-1].replace('Strand', '') 
			if strand not in ['+', '-']: print('\n\t(!) RAMPAGE signal file: Could not identify genomic strand. \n\n', sys.exit()) 
		elif line.startswith('variableStep'):
			chromosome = line.rstrip().split("=")[-1]
			if chromosome not in parsed_rampage_data: parsed_rampage_data[chromosome] = {'+': {}, '-': {}} 
		else: 
			line = line.rstrip().split('\t')
			position = int(line[0]); signal = float(line[1])
			parsed_rampage_data[chromosome][strand][position] = signal
	rampage_file.close()
	## Filter out undesired chromosomes: 
	for chromosome in excluded_chr: 
		if chromosome in parsed_rampage_data: del parsed_rampage_data[chromosome] 
	## Get some stats: 
	total_number_of_rampage_tags = 0
	number_rampage_tags_plus_strand = 0; number_rampage_tags_minus_strand = 0 
	for chromosome in parsed_rampage_data: 
		chr_tag_counts[chromosome] = 0 
		current_count = 0 
		for position in parsed_rampage_data[chromosome]['+']: current_count += parsed_rampage_data[chromosome]['+'][position] 
		number_rampage_tags_plus_strand += current_count 
		chr_tag_counts[chromosome] += current_count 
		current_count = 0 
		for position in parsed_rampage_data[chromosome]['-']: current_count += parsed_rampage_data[chromosome]['-'][position] 
		number_rampage_tags_minus_strand += current_count 
		chr_tag_counts[chromosome] += current_count 
	total_number_of_rampage_tags = number_rampage_tags_plus_strand + number_rampage_tags_minus_strand 
	## Filter out chromosomes without signal: 
	for chromosome in chr_tag_counts: 
		if chr_tag_counts[chromosome] == 0: del parsed_rampage_data[chromosome] 
	## Return: 
	return parsed_rampage_data, total_number_of_rampage_tags, number_rampage_tags_plus_strand, number_rampage_tags_minus_strand, chr_tag_counts 


######################## DEFINE FUNCTION TO LOAD & PARSE Read2 BACKGROUND DATA: 
def load_background_data(read2_background_file): 
	background_file = open(read2_background_file) 
	background_data = {} 
	for line in background_file: 
		if line.startswith('track'): 
			trackLine = line.rstrip().split(' ') 
			strand = None 
			for field in trackLine: 
				if field.startswith('name='): strand = field.split('_')[-1].replace('Strand', '') 
			if strand not in ['+', '-']: print('\n\t(!) Read 2 Background file: Could not identify genomic strand. \n\n', sys.exit()) 
			continue 
		elif line.startswith('variableStep'): 
			chromosome = line.rstrip().split("=")[-1] 
			if chromosome not in background_data: background_data[chromosome] = {'+': {}, '-': {}} 
			continue 
		line = line.rstrip().split('\t') 
		position = int(line[0]); signal = float(line[1]) 
		background_data[chromosome][strand][position] = signal 
	background_file.close() 
	return background_data 


######################## DEFINE FUNCTION TO COMPUTE RAMPAGE TAG DENSITY OF FISRT WINDOW OF CURRENT CHROMOSOME:
def get_rampage_initial_window(chromosome, sliding_window_width, rampage):
    window_start = 1                                                                                                    # Window start coordinate INCLUSIVE. 
    window_end = window_start + sliding_window_width - 1                                                                # Window end coordinate EXCLUSIVE.
    for genome_strand in ['+', '-']:
        rampage_tags = 0
        for individual_position in range(window_start, window_end):                                                     # (!) EXCLUDE LAST POSITION! Will be considered "new position" in first round. 
            rampage_tags_at_position = rampage[genome_strand].get(individual_position, 0)
            rampage_tags += rampage_tags_at_position
        if genome_strand is '+': rampage_tags_plus_strand = rampage_tags
        elif genome_strand is '-': rampage_tags_minus_strand = rampage_tags
    return rampage_tags_plus_strand, rampage_tags_minus_strand 


######################## DEFINE FUNCTION TO COMPUTE ONE-TAILED P-VALUES UNDER NEGATIVE BINOMIAL MODEL: 
def get_negative_binomial_p_value(average_per_position, window_size, dispersion_parameter, number_rampage_tags): 
	average_per_window = average_per_position * window_size 
	success_probability = dispersion_parameter / (average_per_window + dispersion_parameter) 
	p_value = scipy.stats.nbinom.sf(number_rampage_tags, dispersion_parameter, success_probability)
	if math.isnan(p_value): p_value = 0.0
	return p_value


######################## DEFINE FUNCTION TO RUN FDR CORRECTION: 
def fdr_correction(p_values_list, false_discovery_rate): 
    p_values_list.sort()
    k = 0
    m = float(len(p_values_list))
    while ((k<len(p_values_list)) and (p_values_list[k] <= (((k+1)/m) * false_discovery_rate))):                       # List indexing is 0-based, k-indexing for FDR is 1-based. 
        k+=1
    if k == len(p_values_list):
        print("\n(!) FDR correction failure: not enough p-values saved. \n\n")
        sys.exit()
    else: FDR_threshold = k
    i = k-1                                                                             # Check that not many P-values of index > k are the same as P-value(k):
    identical_p_values = 0                                                              # it would lead to too many windows being called.
    while p_values_list[i+1] == p_values_list[i]:
        identical_p_values += 1
        i+=1
    if identical_p_values > 10:                                                         # If many identical p-values: be conservative and pick the one immediately smaller. 
        k=0
        while p_values_list[k] < p_values_list[FDR_threshold]:
            k+=1
        FDR_threshold = k
    p_value_threshold = p_values_list[FDR_threshold] 
    return p_value_threshold


######################## DEFINE FUNCTION TO SORT OUT (AND COUNT) SIGNIFICANT WINDOWS AFTER MULTIPLE TESTING CORRECTION: 
def get_significant_windows(p_values_file_name, p_value_threshold): 
    window_p_values_file = open(p_values_file_name)
    number_significant_windows = 0
    sig_windows = {}
    for strand in ['+', '-']: sig_windows[strand] = {}
    for line in window_p_values_file:
        parsed_line = line.rstrip("\n").split("\t")
        p_value = float(parsed_line[0])
        if p_value < p_value_threshold: 
            strand = parsed_line[1]
            window_start = int(parsed_line[2])
            window_end = int(parsed_line[3])
            rampage_tags_in_window = int(parsed_line[4])
            sig_windows[strand][window_start] = [window_end, rampage_tags_in_window]
            number_significant_windows += 1
    window_p_values_file.close()
    os.remove(p_values_file_name)
    return (sig_windows, number_significant_windows)


######################## DEFINE FUNCTION TO FUSE NEIGHBORING WINDOWS INTO PEAKS: 
def fuse_consecutive_windows(windows_chr_strand, maximum_distance, chromosome_size, rampage):
    rampage_array = numpy.zeros((chromosome_size,), dtype=int)
    for index in range(chromosome_size):
        genomic_position = index + 1
        rampage_signal = rampage.get(genomic_position, 0)
        rampage_array[index] = rampage_signal
    inter_window_distances_list = []
    sorted_coordinates = sorted(windows_chr_strand.keys())
    current_peak_start = sorted_coordinates[0]
    for i in range(len(sorted_coordinates)-1):                                                      # Compute distance between each window (except last) and the next one down the chromosome. 
        current_start_position = sorted_coordinates[i]
        next_start_position = sorted_coordinates[i+1]
        distance = next_start_position - current_start_position                
        inter_window_distances_list.append(distance)
        if distance <= maximum_distance:
            next_end_position = windows_chr_strand[next_start_position][0]
            windows_chr_strand[current_peak_start][0] = next_end_position 
            del windows_chr_strand[next_start_position]
        else:
            current_peak_start = next_start_position
    for start_position in windows_chr_strand:
        end_position = windows_chr_strand[start_position][0] 
        tag_count = numpy.sum(rampage_array[start_position - 1 : end_position])                         # Positions 1-based, array indexes 0-based. INCLUDE end position in window.
        windows_chr_strand[start_position][1] = tag_count 
    return windows_chr_strand, inter_window_distances_list

     
######################## DEFINE FUNCTION TO TRIM PEAKS DOWN TO THE FIRST & LAST POSITIONS WITH RAMPAGE DATA: 
def trim_rampage_peaks(windows_chr_strand, rampage): 
    sorted_coordinates = sorted(windows_chr_strand.keys())
    number_of_rampage_peaks = 0
    for position in sorted_coordinates:
        number_of_rampage_peaks += 1
        peak_data = windows_chr_strand[position]
        peak_start = position; peak_end = peak_data[0]
        while peak_start not in rampage: peak_start += 1
        while peak_end not in rampage: peak_end -= 1
        del windows_chr_strand[position]
        windows_chr_strand[peak_start] = [peak_end, peak_data[1]] 
    del sorted_coordinates
    return windows_chr_strand, number_of_rampage_peaks


######################## DEFINE FUNCTION TO SCAN INDIVIDUAL CHROMOSOMES: 
def scan_chromosome(arguments): 

	start_chromosome_time = time.time()

	## Parse arguments: 
	kro = arguments[0]
	kro_sizes_record = arguments[1]
	tag_count = arguments[2]
	rampage_data_kro = arguments[3]
	sliding_window_size = arguments[4] 
	background_signal = arguments[5] 
	background_weight = arguments[6] 
	threshold_for_p_values = arguments[7] 

	## Exclude chromosomes with no RAMPAGE data: 
	if tag_count == 0:  
		if options.verbose: print("\n%s:\tNo RAMPAGE data" % kro)
		return	
	window_p_values_file_name = temp_file_names_root + "%s_P-values_Wsize%snt.txt" %(kro, sliding_window_size)
	window_p_values_file = open(window_p_values_file_name, "w")

	## Define number of tags of initial window on current chromosome: 
	chromosome_length = kro_sizes_record[kro] 
	rampage_end_windows = get_rampage_initial_window(kro, sliding_window_size, rampage_data_kro)
	rampage_tags_last_window = {} 
	rampage_tags_last_window['+'] = rampage_end_windows[0] 
	rampage_tags_last_window['-'] = rampage_end_windows[1] 
	rampage_tags_departing_position = {}
	rampage_tags_departing_position['+'] = 0 
	rampage_tags_departing_position['-'] = 0
	rampage_tags_new_position = {} 
	
	## Get read 2 background info: 
	bg_plus = 0; bg_minus = 0  
	for k in range(1, sliding_window_size): 
		bg_plus += background_signal['+'].get(k, 0) 
		bg_minus += background_signal['-'].get(k, 0) 
	background_last_window = {'+': bg_plus, '-': bg_minus} 
	background_departing_position = {'+': 0, '-': 0} 
	background_new_position = {} 

	## Slide window (size k) along chromosome:
	first_value_written = False 
	all_p_values = [] 
	start = time.time()
	for position in range(1, (2 + chromosome_length - sliding_window_size)):                                        # Coordinates are 1-based & end of "range" is exclusive.
		
		## Call clusters on chromosome, for each strand:
		window_start = position                                                                                     # Window start coordinate INCLUSIVE. 
		window_end = position + sliding_window_size - 1                                                             # Position is 1-based. End coordinate INCLUSIVE. 
		for strand in genome_strands:

			# Get RAMPAGE signal new window:
			rampage_tags_new_position[strand] = rampage_data_kro[strand].get(window_end, 0)
			rampage_tags_in_window = rampage_tags_last_window[strand] - rampage_tags_departing_position[strand] + rampage_tags_new_position[strand]
			
			# Get background info & Correct signal: 
			background_new_position[strand] = background_signal[strand].get(window_end, 0) 
			background_in_window = background_last_window[strand] - background_departing_position[strand] + background_new_position[strand] 
			background_tags = background_in_window * background_weight 
			corrected_tag_count = int(round(rampage_tags_in_window - background_tags)) 
			rampage_tags_in_window = max(0, corrected_tag_count) 
			
			# Define first position of this window as "departing position" for next window:
			rampage_tags_last_window[strand] = rampage_tags_in_window
			rampage_tags_departing_position[strand] = rampage_data_kro[strand].get(window_start, 0) 
			background_last_window[strand] = background_in_window 
			background_departing_position[strand] = background_signal[strand].get(window_start, 0) 

			# Calculate probability of getting AT LEAST the observed RAMPAGE tag density under null model (One-tailed):
			if rampage_tags_in_window == 0: p_value = 1.0
			else:
				if rampage_tags_in_window <= max_possible_number_rampage_tags: p_value = p_values_table[rampage_tags_in_window] 
				else: p_value = get_negative_binomial_p_value(RAMPAGE_genome_average, sliding_window_size, negative_binomial_dispersion_parameter, rampage_tags_in_window) 
			if not first_value_written: 
				window_p_values_file.write("%s\t%s\t%s\t%s\t%s\n" %(p_value, strand, window_start, window_end, rampage_tags_in_window)) 
				first_value_written = True 
			elif p_value < threshold_for_p_values * FDR:
				window_p_values_file.write("%s\t%s\t%s\t%s\t%s\n" %(p_value, strand, window_start, window_end, rampage_tags_in_window))
				all_p_values.append(p_value)

	window_p_values_file.close()
	finish_chromosome_time = int((time.time() - start_chromosome_time) / 60)
	return all_p_values 


######################## DEFINE FUNCTION TO IDENTIFY & CLUSTER SIGNIFICANT WINDOWS:  
def cluster_significant_windows(arguments): 
	# Parse arguments: 
	chr = arguments[0] 
	pvalues_filename = arguments[1] 
	threshold = arguments[2]
	maxdist = arguments[3]
	chr_size = arguments[4]
	rampage_data = arguments[5]
	verbose = arguments[6] 
	# Identify significant windows: 
	sorted_windows = get_significant_windows(pvalues_filename, threshold)
	significant_windows = sorted_windows[0]
	chr_significant_windows = sorted_windows[1]
	if verbose: 
		print("\nCurrent chromosome: \t\t\t", chr) 
		print("Significant windows: \t\t\t", chr_significant_windows)
	# Compute distances separating consecutive windows and fuse together neighboring windows if conditions satisfied: 
	nb_peaks = 0 
	for strand in ['+', '-']:
		if len(significant_windows[strand]) > 1: 
			fused_windows = fuse_consecutive_windows(significant_windows[strand], maxdist, chr_size, rampage_data[strand])
			significant_windows[strand] = fused_windows[0]
			# For each peak, define boundaries as the first & last positions to have at least 1 RAMPAGE tag:
			trimmed_peaks = trim_rampage_peaks(significant_windows[strand], rampage_data[strand])
			significant_windows[strand] = trimmed_peaks[0]
			nb_peaks += trimmed_peaks[1]
	return (significant_windows, nb_peaks) 


#################################################################################################################################################
#################################################################################################################################################


#### Define command line options & arguments:
parser.add_option("--width", "-w", type=int, action="store", dest="width", default=15)
parser.add_option("--NBinom", "-n", type=float, action="store", dest="NBinom", default=0.6)
parser.add_option("--fdr", "-f", type=float, action="store", dest="fdr", default=0.000001)
parser.add_option("--maxdist", "-d", default=150, type=int, action="store", dest="maxdist")
parser.add_option("--read2-background", type=str) 
parser.add_option("--bg-weight", type=float, default=0.5) 
parser.add_option('--verbose', '-v', action="store_true", default=False) 
parser.add_option('--exclude', '-x', type=str, default='chrM') 
parser.add_option('--threads', '-t', type=int, default=1) 
(options, args) = parser.parse_args() 
if len(args) == 2: 
	chrSizesFileName = args[0] 
	rampage_file_name = args[1] 


#### Process input: 
if len(args) == 2: 
	if not rampage_file_name.startswith('/'): rampage_file_name =  os.getcwd() + '/' + rampage_file_name 
	excluded_chromosomes = options.exclude.split(',') 


#### Print usage message: 

print('''\n
This script takes as input a WIGGLE file describing the density of RAMPAGE read 5' ends 
over the genome. For a sliding window of size k sliding through each position across the 
genome, it computes the probability of obtaining the observed number of RAMPAGE tags under 
the null model. The null model is a negative binomial distribution whose mean is the 
observed average density of tags per window, and whose dispersion parameter is user-defined. 

Optionally, it can also be provided with a second wiggle file describing the density of 
RAMPAGE downstream reads (i.e., the read of the mate pair that starts at the RT priming 
site). The downstream read coverage is used to subtract a pseudocount attributable to 
background signal prior to computing the p-value for each window. The weight to be given 
to this backgroung estimate is a user-defined parameter. 

The p-values are corrected by the method of Benjamini & Hochberg and thresholded to allow 
for an FDR below a value defined by the user. All significant windows within a distance d 
of each other are then merged, which generates the final peaks. 


ARGUMENTS:

\t- Argument 1:\t\tFile providing the size of each chromosome in the reference 
\t\t\t\tgenome (Format <chrName\\tchrSize\\n>). 
\t- Argument 2:\t\tRAMPAGE tag density file (Wiggle). 
\t\t\t\tThe track line for each strand is required to contain a name
\t\t\t\tfield ending with "_+" or "_-", from which the strand is 
\t\t\t\tdeduced. 


OPTIONS:

\t--width (-w)\t\tSliding window width. Default: 15 bases. 
\t--NBinom (-n)\t\tDispersion parameter of the negative binomial. Default: 0.6. 
\t--read2-background\tDownstream read density file (wiggle). 
\t--bg-weight (-w)\tWeight to be given to the background signal. Default: 0.5. 
\t--fdr (-f)\t\tFalse discovery rate. Default: 10^(-6). 
\t--exclude (-x)\t\tComma-separated list of chromosomes to exclude. Default: "chrM". 
\t--distance (-d)\t\tMaximum distance for merging neighboring windows. Default: 150. 
\t--threads (-t)\t\tNumber of threads to be used. Default: 1. 


In our experience, the dispersion parameter (--NBinom) and the FDR (--fdr) are the most critical
ones to change when optimizing the specificity of the peakcalling.  \n\n\n******************\n\n''')

if len(args) != 2: sys.exit() 


#### Get chromosome names & sizes, and list of mappable positions if desired: 
chr_sizes_record = {}
chrSizesFile = open(chrSizesFileName) 
for line in chrSizesFile: 
	line = line.rstrip().split('\t') 
	chr = line[0] 
	if chr == 'phiX174' or chr.startswith('ERCC-'): continue 
	chr_sizes_record[chr] = int(line[1]) 
chrSizesFile.close() 
for chr in list(chr_sizes_record.keys()): 
	if chr in excluded_chromosomes: del chr_sizes_record[chr] 
number_chromosomes = len(list(chr_sizes_record.keys())) 
total_number_positions = sum(chr_sizes_record.values()) 
number_genomic_positions = 2 * total_number_positions 
print("Total number genomic positions: \t%s" % total_number_positions) 


#### Depending on genome, pick appropriate threshold for raw p-values to be saved for FDR correction: 
if number_chromosomes > 1000: p_value_trashing_threshold = 500 
else: p_value_trashing_threshold = 20 


#### Define parameters for peak calling:
FDR = options.fdr
sliding_window_size = options.width
negative_binomial_dispersion_parameter = options.NBinom
max_distance = options.maxdist
if max_distance == 12345678: max_distance = sliding_window_size


#### Get output file names root:
output_file_names_root = rampage_file_name.replace('.wig', '_RAMPAGEpeaks')  
null_model_distribution = "NBinom%.2f" % options.NBinom
temp_directory = os.getcwd() + '/.TEMP_%s/' % time.time()  
os.mkdir(temp_directory) 
temp_file_names_root = temp_directory + output_file_names_root.split('/')[-1] 


#### If desired, load read 2 background data: 
rampage_data_loading_start = time.time()
if options.read2_background: 
	background_filename = options.read2_background 
	print('Read 2 background file: \t\t', background_filename) 
	print('Background signal weight: \t\t', options.bg_weight) 
	print('Loading background data...') 
	options.bg_weight = options.bg_weight / sliding_window_size 
	background = load_background_data(background_filename) 
else: 
	background = {} 
	print('No read 2 background correction.') 


#### Open RAMPAGE Wiggle file & Store data in memory:
print("RAMPAGE data file: \t\t\t", rampage_file_name.split("/")[-1])
print("Loading RAMPAGE data...")
parsed_rampage_data_file = load_rampage_data(rampage_file_name, excluded_chromosomes)
rampage_data = parsed_rampage_data_file[0] 
total_number_of_rampage_tags = int(parsed_rampage_data_file[1])
number_rampage_tags_plus_strand = int(parsed_rampage_data_file[2])
number_rampage_tags_minus_strand = int(parsed_rampage_data_file[3])
chr_tag_counts = parsed_rampage_data_file[4] 
rampage_loading_time = time.time() - rampage_data_loading_start 
print("Loading time: \t\t\t\t%.0f s\n" % rampage_loading_time)
print("Total number of RAMPAGE tags: \t\t", total_number_of_rampage_tags)
print("RAMPAGE tags on + strand: \t\t", number_rampage_tags_plus_strand)
print("RAMPAGE tags on - strand: \t\t", number_rampage_tags_minus_strand)


#### Filter RAMPAGE data (chromosomes not in reference): 
rampage_data_unfiltered = rampage_data 
rampage_data = {} 
for chromosome in rampage_data_unfiltered: 
	if chromosome in chr_sizes_record: rampage_data[chromosome] = rampage_data_unfiltered[chromosome] 
del rampage_data_unfiltered 


#### BACKGROUND: Calculate expected number of RAMPAGE tags per position:
RAMPAGE_genome_average = total_number_of_rampage_tags / number_genomic_positions
print("Genome average RAMPAGE tag density: \t%0.6f tags/base" % RAMPAGE_genome_average)
print("Sliding window size: \t\t\t%s nt" % sliding_window_size)
print("Dispersion parameter: \t\t\t",  negative_binomial_dispersion_parameter) 
print("FDR for peak calling: \t\t\t%.7f%%" % (FDR*100))
print("Max distance for merging: \t\t%s bases" % max_distance)


#### Generate table of precomputed p_values (speed-up):
p_values_table_start = time.time()
print("\nGenerating p-values table...")
p_values_table = {}
max_possible_number_rampage_tags = 5000
for possible_number_rampage_tags in range(1, max_possible_number_rampage_tags+1):
	p_value = get_negative_binomial_p_value(RAMPAGE_genome_average, sliding_window_size, negative_binomial_dispersion_parameter, possible_number_rampage_tags) 
	p_values_table[possible_number_rampage_tags] = p_value 
p_values_table_time = time.time() - p_values_table_start
print("P-values table computation: \t\t%s s" % int(p_values_table_time))


############################################## For each chromosome:
#
#                                                           - Slide window along chromosome
#                                                           - Count RAMPAGE tags in window
#                                                           - Count number of mappable positions in window
#                                                           - Compute probability of observing such a density under null model.
#                                                           - Sort out windows above significance threshold. 


print("\nCalling enriched windows...") 
if options.verbose: print('\n') 
scanStartTime = time.time() 
genome_strands = ['+', '-']
processed_chromosomes = []
pool = multiprocessing.Pool(options.threads)  
all_args = [] 
for chromosome in chr_sizes_record:  
	if chromosome in excluded_chromosomes: continue 
	if chromosome not in rampage_data: continue 
	processed_chromosomes.append(chromosome) 
	background_chr = background.get(chromosome, {'+': {}, '-': {}}) 
	my_args = (chromosome, chr_sizes_record, chr_tag_counts[chromosome], rampage_data[chromosome], sliding_window_size, background_chr, options.bg_weight, p_value_trashing_threshold) 
	all_args.append(my_args) 
print('Number chromosomes: \t\t\t', len(list(chr_sizes_record.keys()))) 
all_p_values = [] 
all_p_values_by_chromosome = pool.map(scan_chromosome, all_args) 
for item in all_p_values_by_chromosome: all_p_values += item  
del background 
scanTime = (time.time() - scanStartTime) / 60.0 
print('Genome-wide scan: \t\t\t%.0f mins' % scanTime) 


######## Proceed to FDR correction & call significant windows:
FDR_start_time = time.time()
print("\nRunning FDR correction...")
significance_threshold = fdr_correction(all_p_values, FDR)
finish_FDR_time = int(time.time() - FDR_start_time) 
print("Significance threshold: \t\t%.7f" % significance_threshold)
print("Computing time for FDR correction: \t%s s" % finish_FDR_time)


######## For each chromosome, get significant windows & merge window clusters them into peaks:
start_clustering_time = time.time()
print("\nClustering enriched windows...")
number_of_peaks = 0
pool = multiprocessing.Pool(options.threads)  
all_args = [] 
for chromosome in processed_chromosomes:  
	window_p_values_file_name = temp_file_names_root + "%s_P-values_Wsize%snt.txt" %(chromosome, sliding_window_size)
	my_args = (chromosome, window_p_values_file_name, significance_threshold, max_distance, chr_sizes_record[chromosome], rampage_data[chromosome], options.verbose) 
	all_args.append(my_args) 
final_peaks = pool.map(cluster_significant_windows, all_args) 
significant_windows = {}
for k in range(len(processed_chromosomes)): 
	significant_windows[processed_chromosomes[k]] = final_peaks[k][0] 
	number_of_peaks += final_peaks[k][1] 
os.rmdir(temp_directory) 
print("\n\nNumber of RAMPAGE peaks: \t\t%s" % number_of_peaks)
finish_clustering_time = int((time.time() - start_clustering_time) / 60)
print("Clustering time: \t\t\t%s min" % finish_clustering_time)


######## Output list of called peaks to BED6 file: 
RAMPAGE_peaks_file_name = output_file_names_root + '.bed'  
RAMPAGE_peaks_file = open(RAMPAGE_peaks_file_name, "w") 
peak_index = 1 
for chromosome in sorted(significant_windows.keys()): 
	for strand in sorted(significant_windows[chromosome].keys()): 
		for peak_start in sorted(significant_windows[chromosome][strand].keys()): 
			peak_end = significant_windows[chromosome][strand][peak_start][0] 
			rampage_tag_count = significant_windows[chromosome][strand][peak_start][1] 
			peak_start = str(peak_start - 1)  
			peak_end = str(peak_end) 
			peak_ID = 'RAMPAGEpeak_%06.0f' % peak_index 
			BED_line = '\t'.join([chromosome, peak_start, peak_end, peak_ID, str(rampage_tag_count), strand]) + '\n' 
			RAMPAGE_peaks_file.write(BED_line) 
			peak_index += 1 
RAMPAGE_peaks_file.close() 


######## Report summary stats:
if options.verbose: 
	processed_chromosomes.sort() 
	print("\nProcessed chromosomes: \t\t\t", processed_chromosomes[0])
	for item in processed_chromosomes[1:]: print("\t\t\t\t\t", item)
print("\nTotal running time: \t\t\t%s min\n\n" % int((time.time() - start_time) / 60))
```
