## Supplemental_Figures for "Transcription initiation mapping in 31 bovine tissues reveals complex promoter activity, pervasive transcription, and tissue-specific promoter usage"

**TABLE OF CONTENTS**

Supplemental Fig 1 ---------------------------------------------------------------------------------- 2

Supplemental Fig 2 ---------------------------------------------------------------------------------- 3

Supplemental Fig 3 ---------------------------------------------------------------------------------- 4

Supplemental Fig 4 ---------------------------------------------------------------------------------- 6

Supplemental Fig 5 ---------------------------------------------------------------------------------- 7

Supplemental Fig 6 ---------------------------------------------------------------------------------- 8

Supplemental Fig 7 ---------------------------------------------------------------------------------- 9

Supplemental Fig 8 ---------------------------------------------------------------------------------- 9

Supplemental Fig 9 --------------------------------------------------------------------------------- 10

Supplemental Fig 10 ------------------------------------------------------------------------------ 10

Supplemental Fig 11 ------------------------------------------------------------------------------ 11

Supplemental Fig 12 ------------------------------------------------------------------------------ 12

Supplemental Fig 13 ------------------------------------------------------------------------------ 13

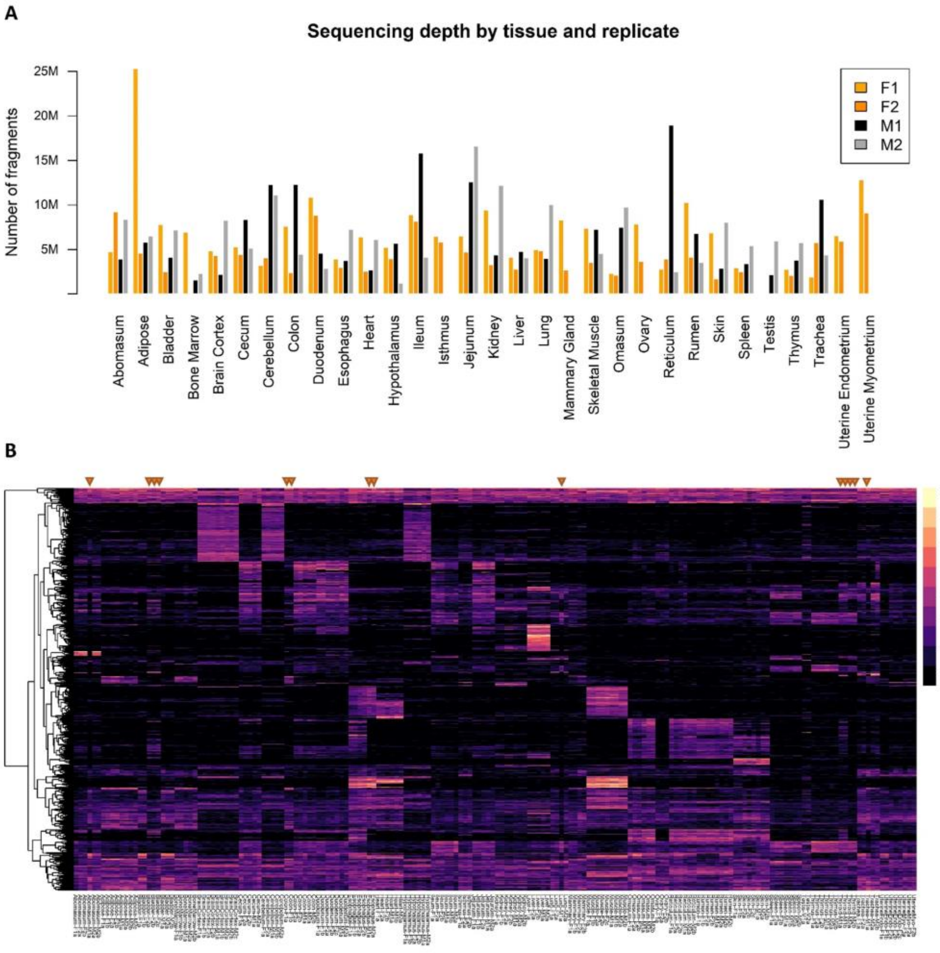


**Supplemental Fig S1.** RAMPAGE dataset composition. (A) Sequencing depth by tissue and replicate. The y axis indicates the number of uniquely mapped fragments after duplicate removal. (B) Profiling by conventional gene counts (variance-stabilized) showing the unclear identity of the abomasum-M1, bladder-F2, colon-F1, esophagus-M1, esophagus-M2, lung-M1, and trachea-M1 samples (indicated with orange triangles), which were excluded from further analyses.

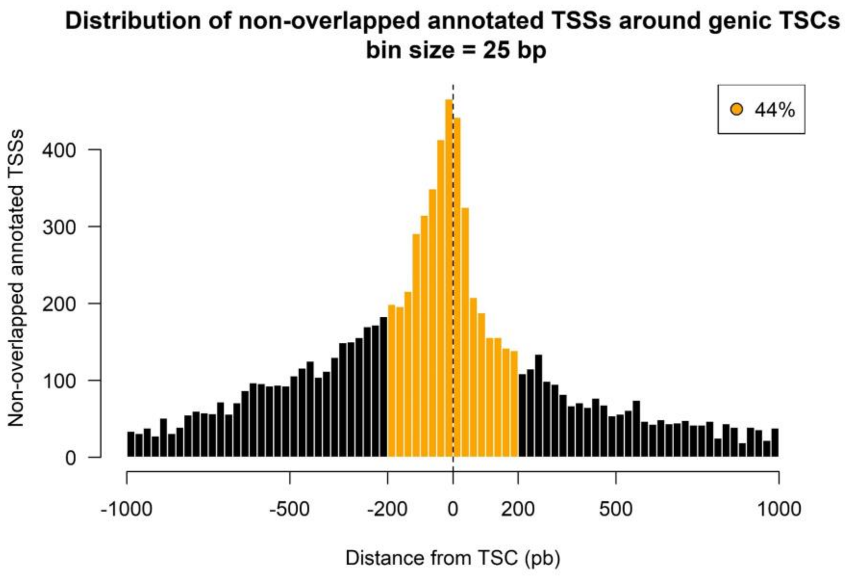


**Supplemental Fig S2.** Distribution of nonoverlapped annotated TSSs relative to genic TSCs identified by RAMPAGE. Many of the annotated TSSs were shifted with respect to TSCs by a margin of 200 bp (orange), while the rest likely belonged to transcripts variants undetected in our dataset (black). Only annotated genes with high expression levels (CPM > 3) were considered for this figure.

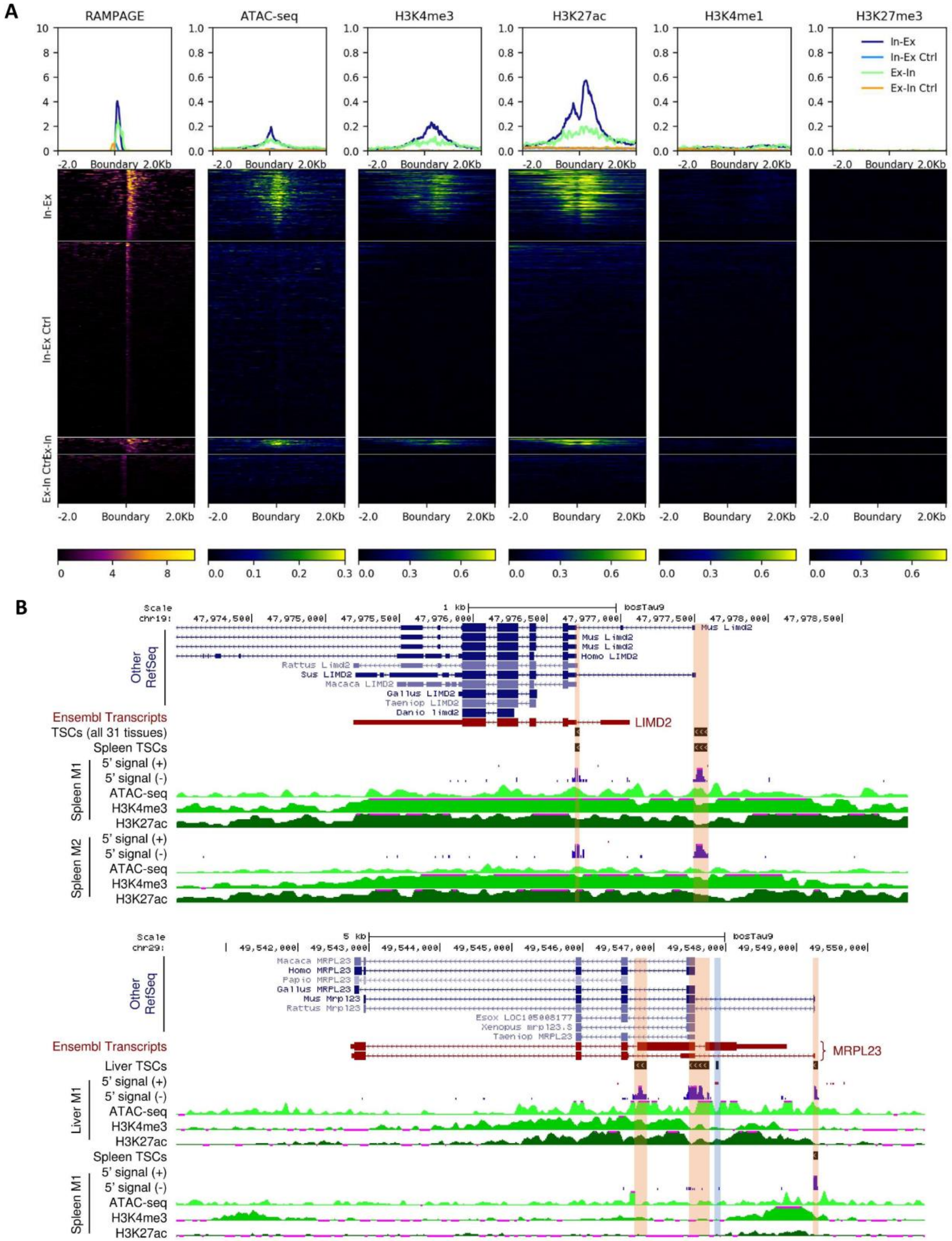


**Supplemental Fig S3.** Chromatin accessibility and histone marks at intron-exon and exon-intron boundaries associated with TSCs (Lung-M2). (A) Distribution of reads from different assays over a window of 2 kb to each side of the boundaries. The RAMPAGE signal contemplates only the first read of the pair (R1). Panels 'In-Ex' and 'Ex-In' represent intron-exon and exon-intron boundaries overlapped by TSCs and expressed at a minimum level of 3 CPM with respect to all RAMPAGE tags. Control regions (Ctrl) comprised boundaries from the same transcripts, covered by at least 3 fragments, and located at least 2 kb away from the nearest TSC to avoid signals from near TSCs. Heatmaps are colored according to CPM values. Profiles above heatmaps represent the median signal at each coordinate of the 4kb window. The co-occurrence of chromatin accessibility (ATAC-Seq) and transcriptional activation marks (H3K4me3, H3K27ac), and the absence of enhancer (H3K4me1) and repressive (H3K27me3) marks at the TSC sites implicated these boundaries as putative promoters. The remaining tissue samples are available as supplementary material. (B) 5' tags and epigenetic signals at boundary-associated TSCs. We found no evidence supporting the annotated TSS for the bovine *LIMD2* gene. In addition, one of the TSCs overlapped an annotated intron-exon boundary. In case of the *MRPL23* gene, we found a highly active TSC at an exon-intron boundary, a second TSC between an exon-intron and an intron-exon boundary, and a third TSC supporting one of the annotated TSSs.

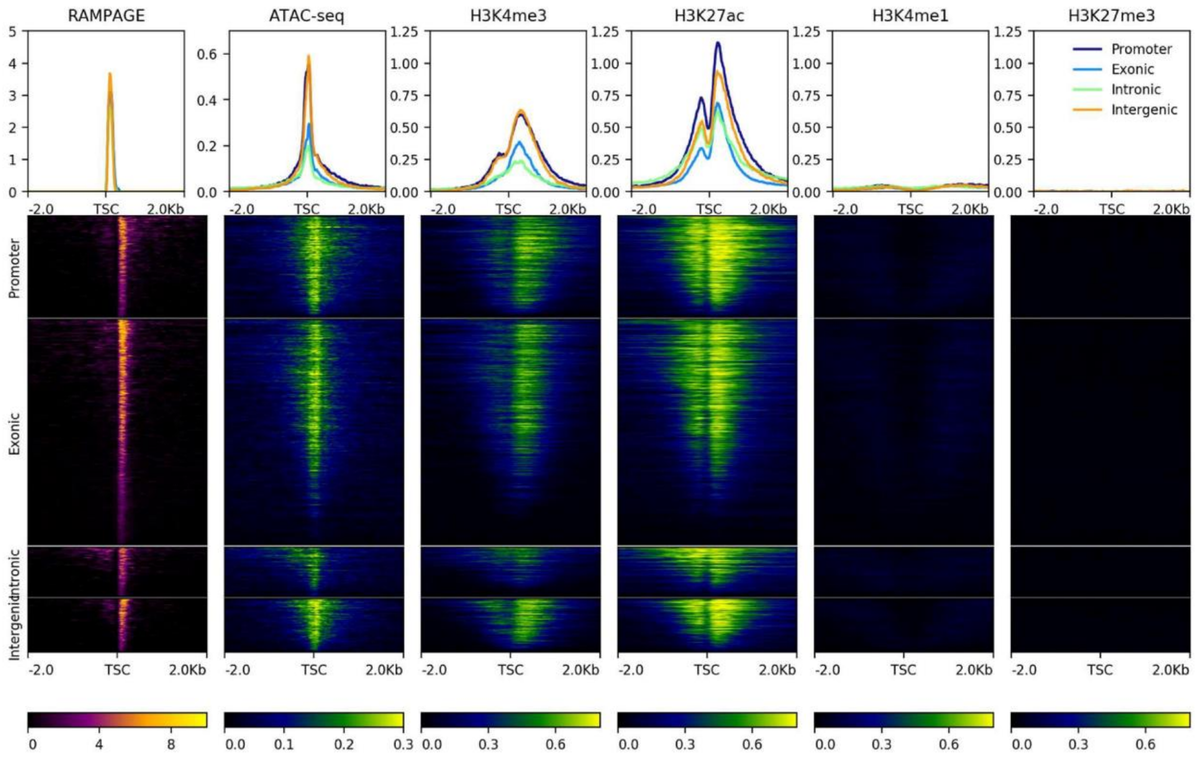


**Supplemental Fig S4.** Epigenetic marks at subgroups of novel genic TSCs defined by their gene location (exon, intron, intergenic region, upstream of annotated TSS (promoter)) in the lung-M2 sample. A consistent co-occurrence of chromatin accessibility and transcriptional activation marks, as well as absence of poised enhancer and repressive marks, was observed in all subgroups. Heatmaps are colored according to CPM values.

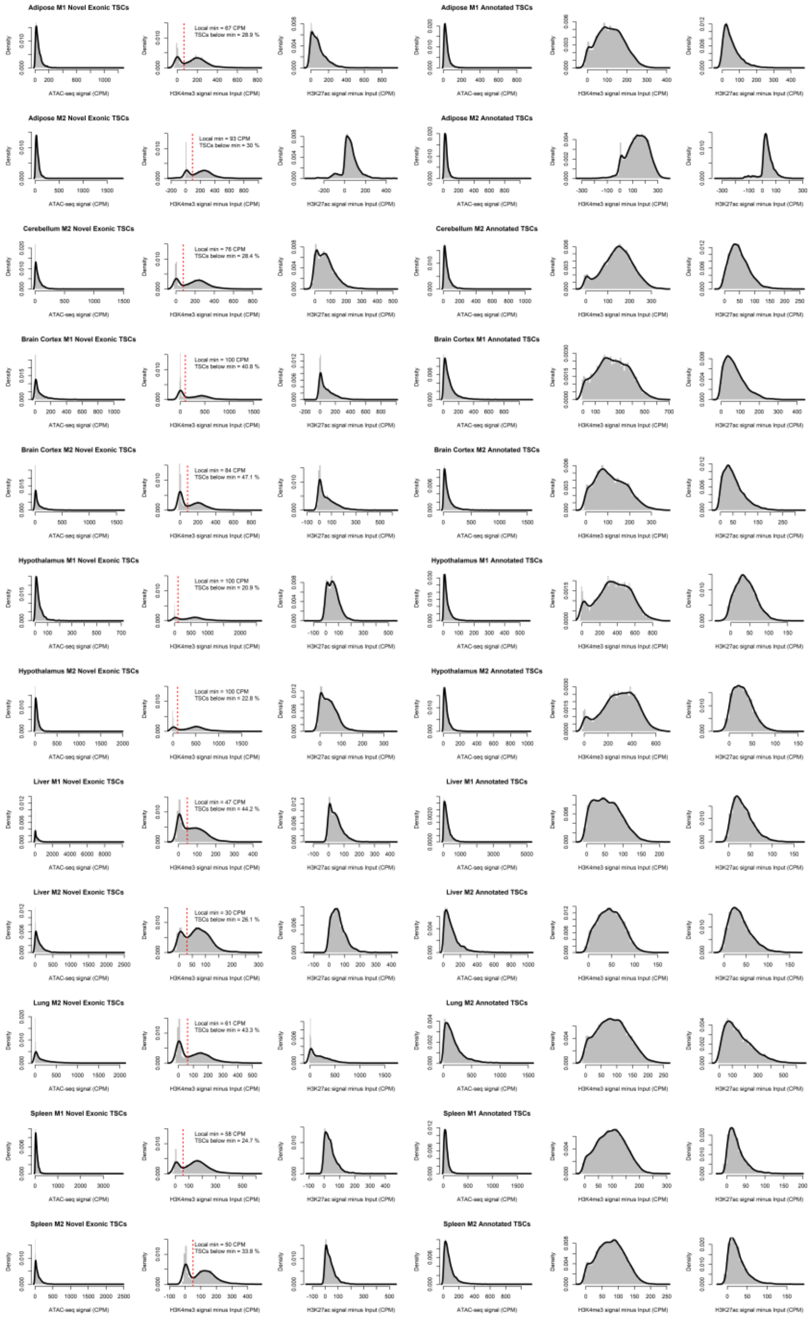

**Supplemental Fig S5**. ATAC-Seq, H3K4me3, and H3K27ac marks around novel TSCs in annotated exons (left) and known TSSs (right). The bimodal distribution of H3K4me3 signal allowed us to estimate that approximately 30% of the novel TSCs in annotated exons could be artifacts produced by incomplete cDNA synthesis or RNA degradation. TSCs under the local minimum of H3K4me3 signal in all samples were labeled in Table S2.

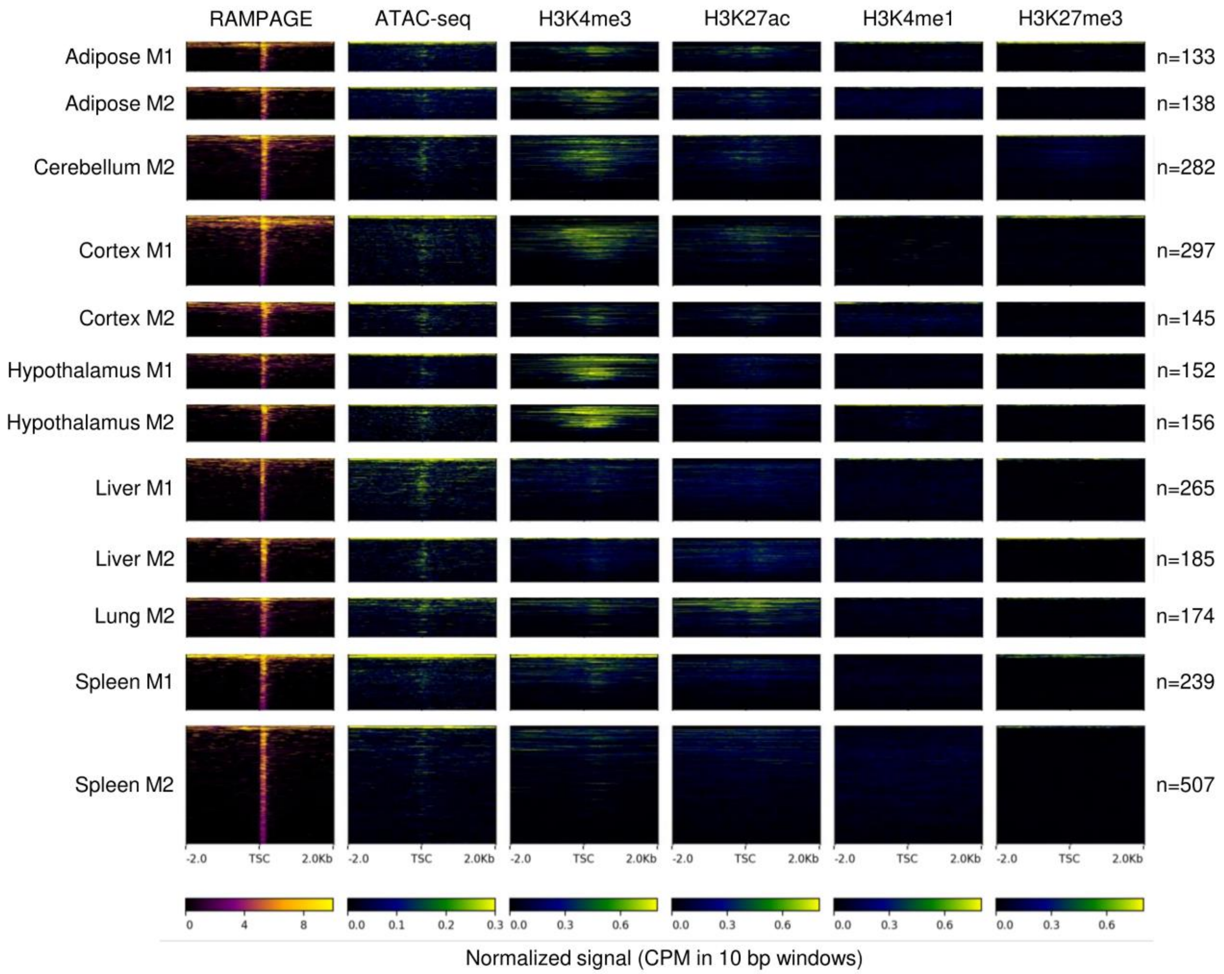


**Supplemental Fig S6**. Epigenetic marks at the remaining unassigned TSCs after prediction of mRNAs, lncRNAs, and pre-miRNAs. From the unclear patterns, we concluded these sites comprised a heterogeneous population of RNAs with unclear functions and technical noise.

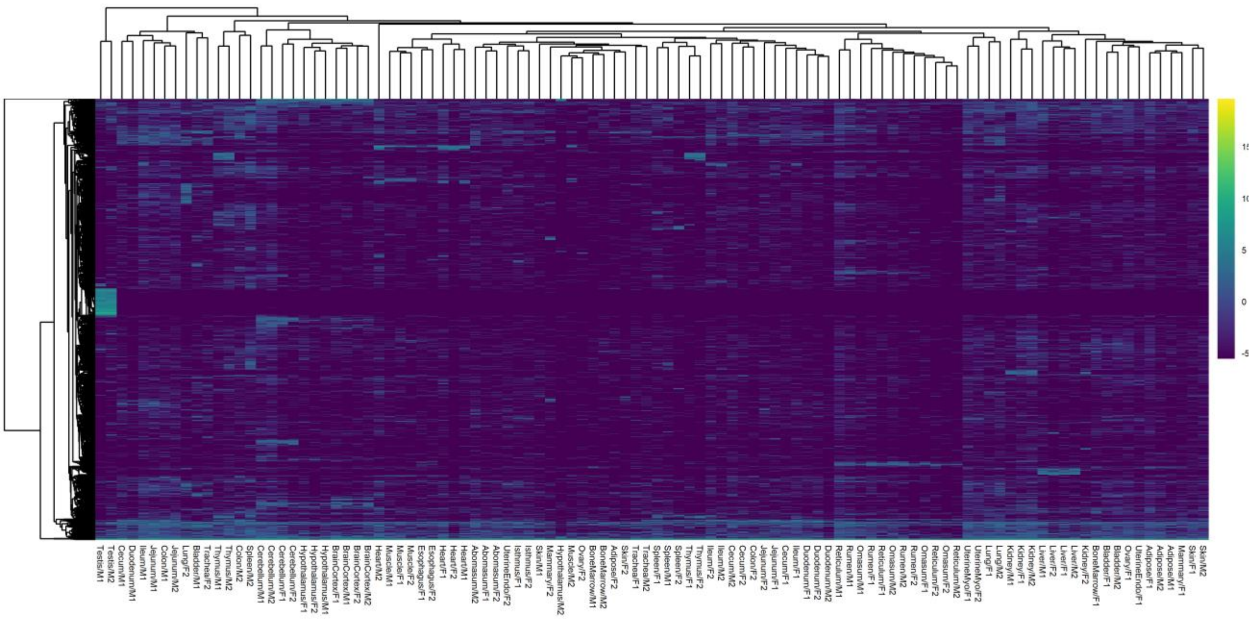

**Supplemental Fig S7.** Hierarchical clustering based on RAMPAGE signal (variance-stabilized counts) at the remaining unassigned TSCs. Samples failed to cluster appropriately, suggesting many of these elements could represent artifacts.

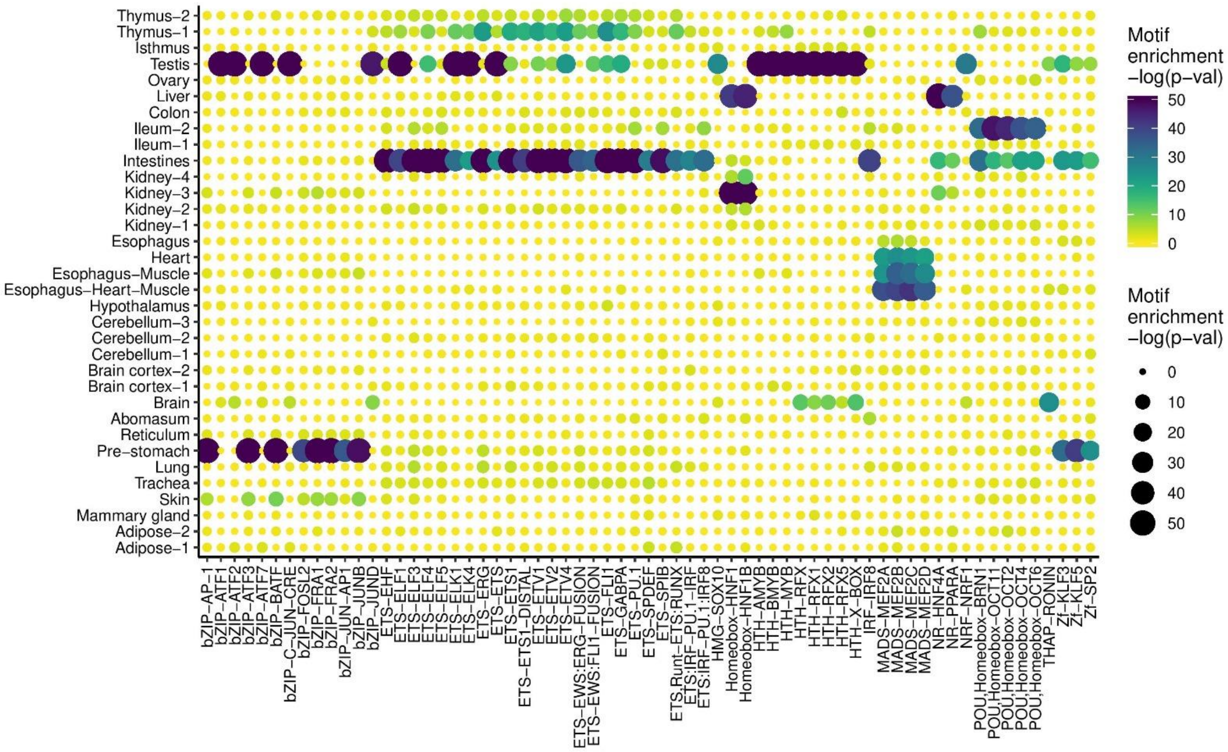


**Supplemental Fig S8**. Binding motif enrichment in regions from 300 bp upstream to 100 bp downstream of co-expressed TSCs.

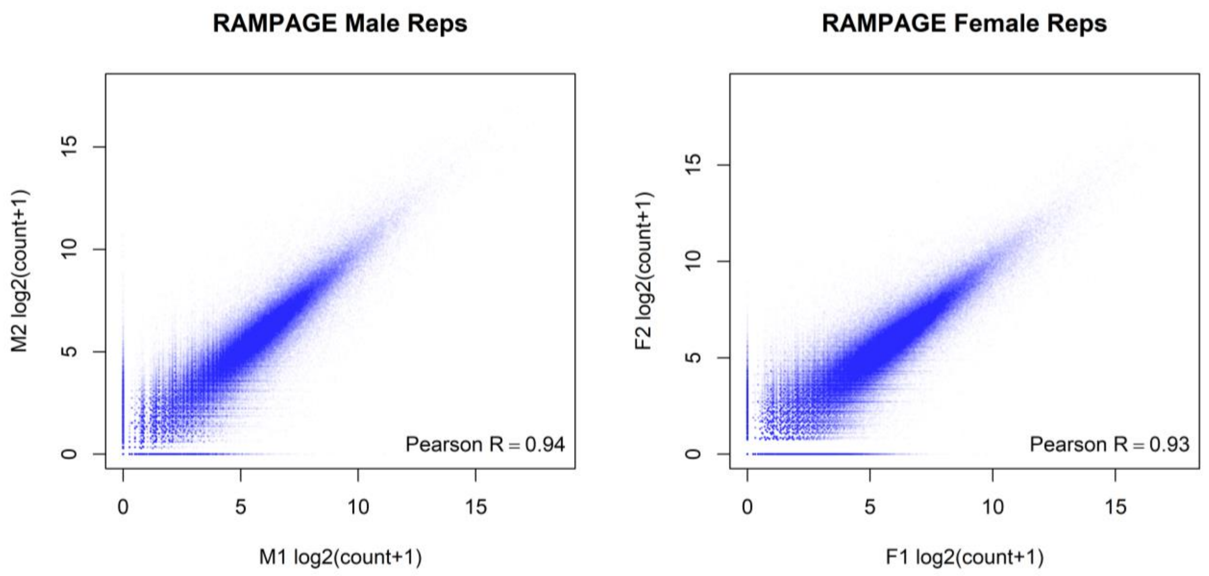


**Supplemental Fig S9.** Reproducibility between RAMPAGE replicates. Counts were normalized by DESEQ2.

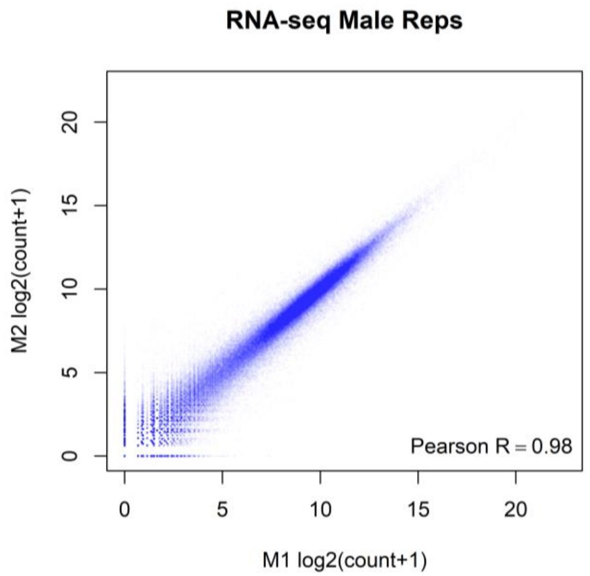


**Supplemental Fig S10**. Reproducibility between replicates for RNA-Seq counts. Counts were normalized by DESEQ2.

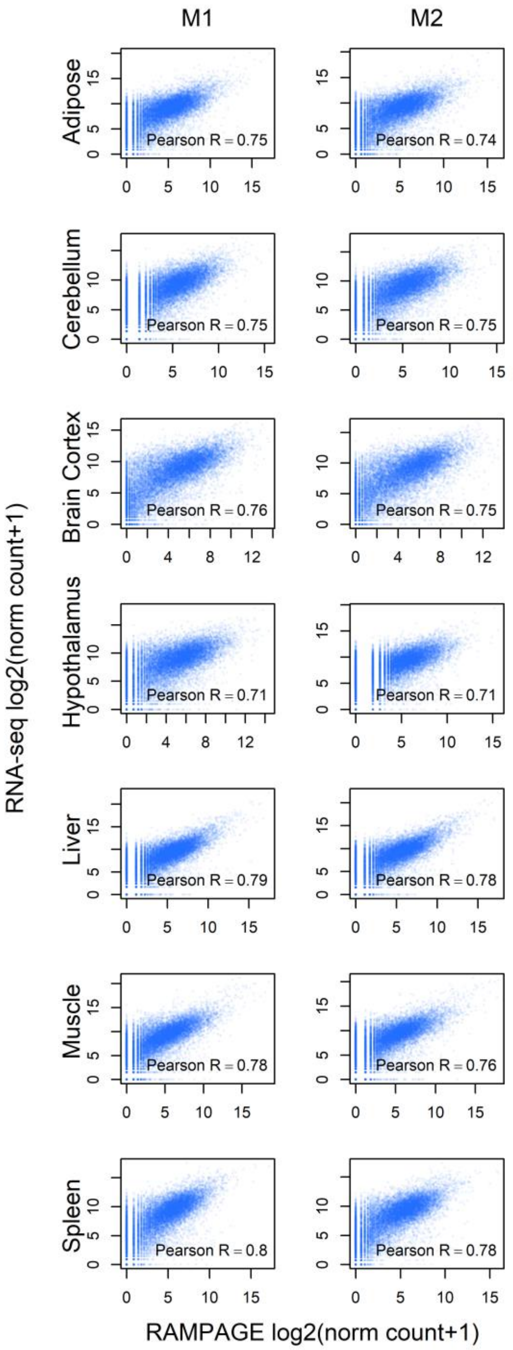


**Supplemental Fig S11**. Pairwise correlations between RNA-Seq and RAMPAGE log-transformed normalized counts. Moderately high correlations were consistently observed across samples. Counts were normalized by DESEQ2.

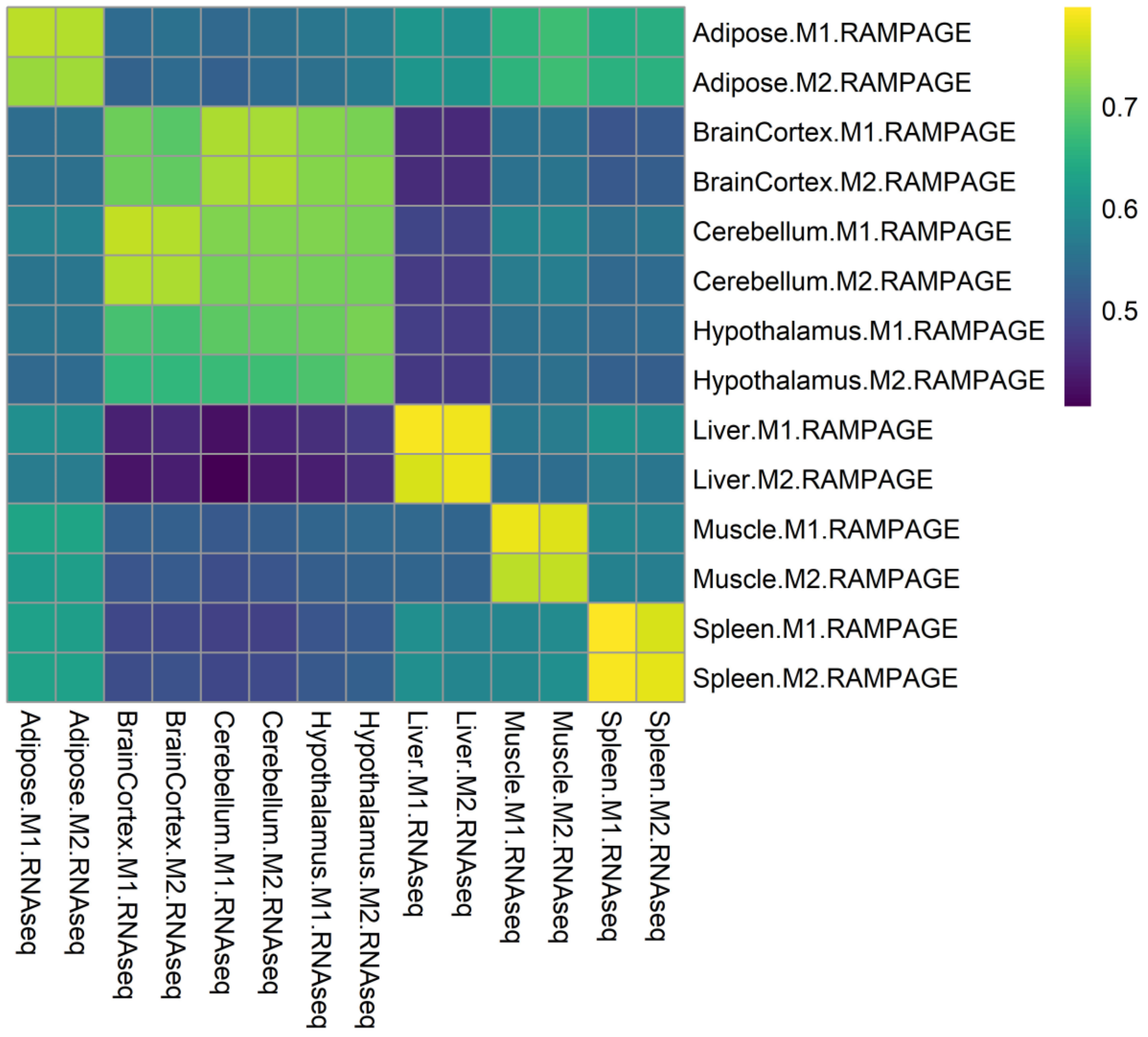


**Supplemental Fig S12**. Pearson correlation between RNA-Seq and RAMPAGE log-transformed normalized counts. High correlation was observed between samples from the same tissue. Counts were normalized by DESEQ2.

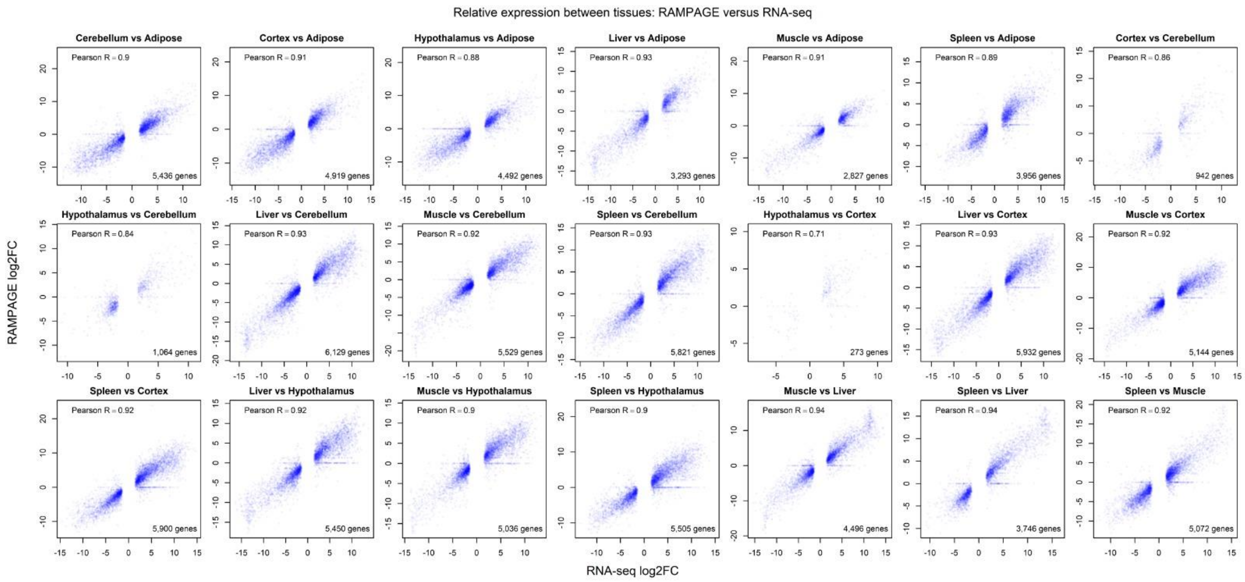


**Supplemental Fig S13.** Pairwise correlations between logFC values obtained from RNA-Seq and RAMPAGE gene counts. The analysis contemplates differentially expressed genes first identified by RNA-Seq. Correlations were high except for brain tissues, where differences in gene expression were minimal.
